# Nuclear receptor LRH-1 regulates early T cell development in mice

**DOI:** 10.64898/2026.05.11.724315

**Authors:** Alice Wiedmann, Nele Käter, Rayan Hassan M. Elshikhidriss, Lea Dietrich, Verena M. Merk, Franziska Rudolf, Daniel F. Legler, Thomas Brunner

## Abstract

T cell development in the thymus requires tightly coordinated transcriptional programs that regulate lineage commitment, proliferation and differentiation. While key transcription factors controlling these processes have been extensively characterized, the contribution of the low expressed nuclear receptor Liver Receptor Homolog 1 (LRH-1, *Nr5a2*) in T cell development remains unexplored. Here, we investigated the role of LRH-1 in thymocyte maturation using an inducible *ex vivo* deletion system and *in vivo* Lck-Cre- and CD4-Cre-mediated LRH-1 knockout mouse models. We demonstrate that inducible LRH-1 deletion impairs early thymocyte development, identifying LRH-1 as a critical regulator of the double negative (DN)2/DN3 to DN4 transition. Early Lck-Cre-mediated deletion of LRH-1, but not CD4-Cre-mediated deletion at the double positive stage, resulted in markedly reduced thymic size and cellularity, indicating a stage-specific requirement for LRH-1 during thymopoiesis. Lck-Cre-mediated LRH-1 deletion led to a decreased frequency of mature CD4⁺ T cells in peripheral lymphoid organs, while the remaining mature T cells were predominantly Cre reporter-negative and therefore escaped LRH-1 deletion. CD4⁺ T cells that escaped Cre-mediated LRH-1 deletion exhibited impaired T cell activation marker expression and cytokine secretion. *In vivo*, these defects resulted in attenuated T cell effector function and compromised regulatory T cell-mediated protection in a T cell transfer model of colitis, indicating impaired effector and regulatory T cell function under (patho)physiological conditions.

Collectively, our findings identify LRH-1 as a critical, previously unrecognized regulator of early thymocyte development, and establish its essential role in shaping functional peripheral CD4⁺ T cell-mediated immune responses.

## Introduction

T cells are the only hematopoietic cells that do not mature in a specific niche in the bone marrow, but in a separate primary lymphoid organ, the thymus. Therefore, common lymphoid progenitors, derived from hematopoietic stem cells, migrate from the bone marrow to the thymus, where they differentiate into early thymic progenitors [1, 2]. These double negative (DN) early thymocytes do not express CD4 or CD8, nor the T cell receptor (TCR), but can be defined by their CD44 and CD25 expression. DN1 cells only express CD44 (CD44^+^ CD25^-^). With the onset of CD25 expression, these cells are classified as DN2 (CD44^+^ CD25^+^) [2, 3]. At these stages, DN1 and early DN2 cells are not yet restricted to T cell lineage development and can still differentiate into other hematopoietic cells, such as B cells, natural killer (NK) cells, innate common lymphoid cells (ILC), dendritic cells and granulocytes. T cell lineage commitment is initially achieved by thymic stromal cell-induced Notch receptor signaling and is irreversible at the end of the DN2 stage [4]. TCRβ rearrangement occurs at DN3 (CD44^lo^ CD25^+^) and DN4 (CD44^-^ CD25^-^), and once successful, massive expansion of thymocytes is accompanied by the upregulation of CD4 and CD8 coreceptors and rearrangement of the TCRα-chain results in double positive (DP, CD4^+^ CD8^+^) T cells. After positive and negative selection, CD4 single positive (SP) or CD8 SP T cells are generated, which egress from the thymus and migrate to peripheral secondary lymphoid organs to exert their function within the immune system [2, 3, 5].

During T cell maturation and T cell lineage commitment, a number of essential transcription factors are involved, such as the T cell factor 1 (TCF-1), GATA binding protein 3 (GATA3), B cell leukemia/lymphoma 11B (BCL11B), Ikaros Zinc Finger family (IkZF) and Runt-related transcription factors (Runx) family members [4]. These key factors not only drive DN cells towards T cell lineage commitment, but also actively suppress the development of other hematopoietic cells. Despite T cell lineage commitment, proliferation in the DN and expansion in the DP stages is obligatory for the development of functional T cells [4, 6].

Another essential transcription factor, whose role in T cell development has not been defined yet, is the nuclear receptor Liver Receptor Homolog 1 (LRH-1, *Nr5a2*). LRH-1 is known to be involved in the regulation of the cell cycle and proliferation, despite other pleotropic organ- and life stage-specific functions [7]. It is mainly expressed in tissues derived from the endoderm but is also present at low levels in immature and mature T cells. Despite its low abundancy, LRH-1 has been shown to be essential for the regulation of cell proliferation in mature murine T cells and human leukemic T cells [8, 9]. Our previous work, employing CD4-Cre-mediated LRH-1 deletion, revealed little effect of LRH-1 deficiency on thymocyte expansion at the DN and DP stage, but demonstrated that LRH-1 is essential for the development of mature functional regulatory and effector T cells [8]. Yet, the precise role of LRH-1 in early T cell development still remains unclear.

In this study, we report for the first time the essential role of LRH-1 in early T cell development using an *ex vivo* inducible LRH-1 deletion system and *in vivo* Lck-Cre and CD4-Cre driven LRH-1 deletion mouse models. Despite its minimal expression in DN thymocytes, the loss of LRH-1 appears to impede thymocyte progression beyond the DN3 stage, but exerting only a negligible effect on DP thymocytes, which exhibit the highest level of LRH-1 expression. Consequently, LRH-1 appears to be critical for the transcriptional control of early but not late thymic T cell development. Importantly, maintenance of LRH-1 expression during early thymocyte development appears to be crucial as only thymocytes escaping LRH-1 deletion can proceed to the mature T cell stage. The resulting mature T cells exert severe defects in their effector functions. These findings highlight the so far unknown essential role of LRH-1 regulated processes in thymic development and T cell-regulated immune responses.

## Results

### Inducible deletion of LRH-1 impairs thymocyte development at the DN3 and DN4 stage

Previous data on human tissues indicate approximately 10- to 100-fold lower *Nr5a2* expression in primary and secondary lymphoid tissues compared to endoderm-derived tissues, such as the liver [10]. Our data show that although this difference is apparently less pronounced in mice, there is still an approximately 2- to 3-fold reduction in *Nr5a2* expression in spleen, splenocytes and thymocytes compared to whole liver lysates (Fig. 1A). A closer look on LRH-1 expression at different stages of T cell development, as defined by the expression of CD44, CD25, CD4 and CD8 [3], showed the highest levels in DP and CD4 SP thymocytes (Fig. 1B). DN1 (CD44^+^ CD25^-^) cells showed the lowest or no detectable expression of LRH-1, whereas its expression increases at the DN2 stage (CD44^+^ CD25^+^) and in some DN3 (CD44^lo^ CD25^+^) cells followed by a recurrent decrease in DN4 (CD44^-^ CD25^-^) thymocytes (Fig. 1B). Based on our observation of differential *Nr5a2* expression in immature T cells, we first established an inducible LRH-1 knockout (iKO) system, in which LRH-1 deletion is achieved *ex vivo* by 4-hydroxytamoxifen (4-HT) treatment. To this end, isolated thymocytes were co-cultured with the murine bone marrow-derived stromal cell line OP9-DL1, expressing the Notch ligand Delta [11], and the presence of murine IL-7 to mimic the thymic niche (Fig. 1C, Suppl. Fig. 1A). The successful activation of Cre recombinase in the iKO mouse model was measured by the membrane-targeted tandem dimer Tomato membrane-targeted GFP (mTmG) reporter construct (Suppl. Fig. 1B) [12]. The induction of Cre recombinase activity after 4-HT treatment and thus successful GFP reporter expression was achieved in about 30% of living thymocytes (Suppl. Fig. 1B). The deletion of LRH-1 led to a significantly decreased *Nr5a2* expression (Fig. 1D) and changes in the frequency of DN1, DN3 and DN4 thymocytes were observed (Fig. 1E). The induction of LRH-1 deletion resulted in an accumulation of DN1 and a strongly reduced frequency of DN3 and DN4, indicating an important role of LRH-1 in early thymocyte development (Fig. 1E). Cre-mediated or 4-HT toxicity could be excluded as no significant changes in the frequency of the different thymocyte subsets were observed in 4-HT-treated LRH-1^L2/L2^ and Rosa26-Cre-ERT2 control mice (Suppl. Fig. 1C-D). Interestingly, up to 80% of DN3 and 60% of DN4 thymocytes were identified as GFP^+^ and thus LRH-1-deficient, whereas deletion was only successful in up to 50% of DN2 and approximately 20% of DN1, DP, CD4 SP and CD8 SP thymocytes (Fig. 1F). Higher concentrations of 4-HT did not further increase Cre recombinase activity visualized by GFP^+^ cells but resulted in higher cell death in 4-HT-treated iKO and Rosa26-Cre-ERT2 control thymocytes (Suppl. Fig. 1E). These findings indicate that the presence of LRH-1 is critical to progress beyond the DN4 stage during thymic development.

**Figure 1:**
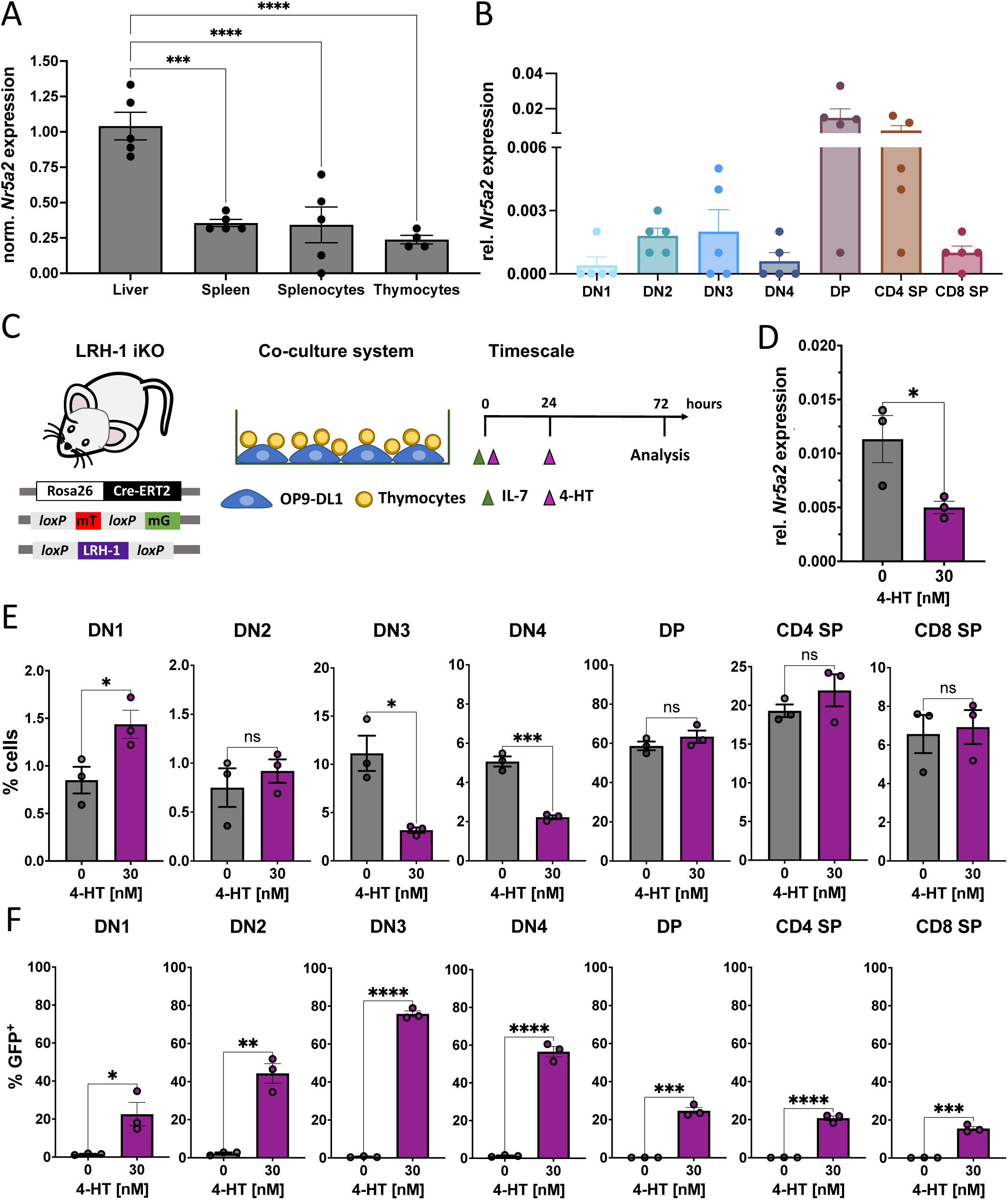
Inducible deletion of LRH-1 impairs thymocyte development at the DN3 and DN4 stage. (A) *Nr5a2* RT-qPCR of liver, spleen, isolated splenocytes and thymocytes sample normalized to *Nr5a2* expression in liver and (B) relative *Nr5a2* RT-qPCR of sorted thymocyte stages from B6 wildtype mice. Gene expression was normalized to *Actb*. Bars show mean ± SEM, dots represent individual mice (n = 4-5). (C) Schematic representation of the LRH-1 iKO (Rosa26-Cre-ERT2 x mTmG x LRH-1, LRH-1 iKO) mouse model used in the *ex vivo* OP9-DL1 and thymocyte co-culture system. For LRH-1 deletion, thymocytes from LRH-1 iKO mice were isolated and treated with IL-7 (1 ng/ml) at t = 0 h and twice with 4-hydroxytamoxifen (4-HT; 30 nM) at t = 0 and t = 24 h after isolation. (D) Relative *Nr5a2* RT-qPCR of isolated LRH-1 iKO thymocytes, either untreated (0 nM) or treated with 4-HT (30 nM) for 24 h. Gene expression was normalized to *Actb*. Bars show mean ± SEM, dots represent individual mice (n = 3 per group). Flow cytometry analysis of the frequency of live thymocyte stages (E) and GFP-positive live thymocyte stages (F) in untreated or twice 30 nM 4-HT treated LRH-1 iKO cells analyzed after 72 h. Bars show mean ± SEM, dots represent individual mice (n = 3 per group). Statistical significance was determined by one-way ordinary ANOVA with Dunnett’s multiple comparison test (A) and unpaired Student’s T-test (D-F), ns: not significant, * p<0.05, ** p<0.005, *** p<0.001, **** p<0.0001. DN = double negative, DP = double positive, SP = single positive.

### LRH-1 deletion in early DN stages, but not in the DP stage results in reduced thymic size and cellularity

Our previous study elucidated that the CD4-Cre-mediated deletion of LRH-1 at the DP stage of T cell development has minor effects on the frequency of immature T cells, but a major impact on the number and effector functions of mature T cells [8]. To further investigate the role of LRH-1 in thymocyte development and to overcome the incomplete deletion of LRH-1 in the 4-HT iKO system, a conditional knockout (cKO) mouse model was used, in which Lck promoter-driven Cre recombinase controls LRH-1 deletion at early DN stages (Lck-Cre cKO). Expression of Cre recombinase under the Lck promoter results in low-level target gene deletion in a subset of DN1 cells, but the Cre activity becomes predominantly evident in DN2 and DN3 thymocytes [13]. These mice were compared to those with CD4 promoter-driven Cre expression and LRH-1 deletion at the DP stage (CD4-Cre cKO) (Fig. 2A) [14].

**Figure 2:**
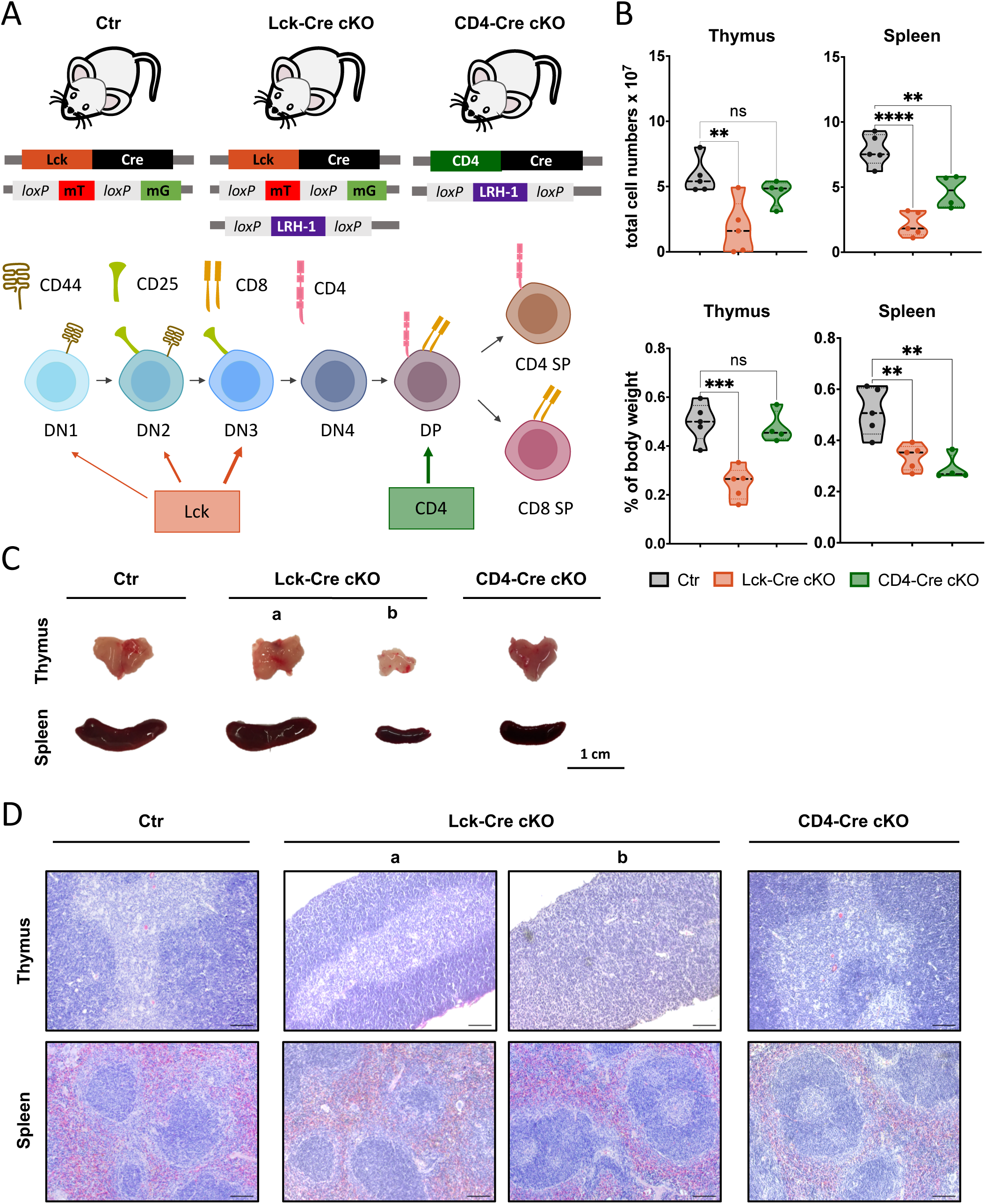
LRH-1 deletion in early DN stages, but not in the DP stage, results in reduced thymic size and cellularity. (A) Schematic showing mouse models for control (Lck-Cre x mTmG; Ctr), LRH-1 cKO in early DN thymocyte stages (Lck-Cre x mTmG x LRH-1; Lck-Cre cKO) and LRH-1 cKO in late DP thymocyte stage (CD4-Cre x LRH-1; CD4-Cre cKO). Partial Lck expression starts at the DN1 and DN2 stage, but occurs at latest at the DN3 thymocyte stage, whereas CD4 expression begins at the DP thymocyte stage. (B) Total thymocyte and splenocyte cell counts (top) and weight of spleen and thymus relative to mouse body weight (bottom) of 9- and 13-week-old Lck-Cre cKO, CD4-Cre cKO and Ctr mice. Each point in the violin plot represents an individual mouse (n = 4-5 per group). (C) Representative images of thymus and spleen of 13-week-old Ctr, Lck-Cre cKO and CD4-Cre cKO mice. (D) Representative H&E-stained sections of thymus and spleen from Ctr, Lck-Cre cKO (a and b) and CD4-Cre cKO mice. Scale bar = 100 µm. Statistical significance was determined by one-way ANOVA with Dunnett’s multiple comparison test, ns: not significant, ** p<0.005, *** p<0.001, **** p<0.0001.

In line with our previous study [8], we observed that CD4-Cre cKO mice had a normal thymus, but a reduced spleen size compared to those of control mice (Fig. 2C; Lck-Cre x mTmG; Ctr). Overall, Lck-Cre- but not CD4-Cre-mediated LRH-1 deletion caused significantly reduced thymus weight and cellularity, whereas spleen weight and cellularity were reduced in both cKO mouse models (Fig. 2B). Surprisingly, Lck-Cre-mediated LRH-1 deletion resulted in two different phenotypes. While some mice had a drastically reduced thymus and spleen size, other mice showed a normal thymus and spleen size (Fig. 2C, a vs. b). Histological analysis revealed that Lck-Cre cKO mice (Fig. 2D; Lck-Cre cKO b) with small thymi lacked a proper separation of thymic medulla and cortex, as seen in Ctr mice. In addition, these mice showed an altered splenic structure with less cellular density in the white pulp, similar to that in CD4-Cre cKO mice, indicating a lower abundance of lymphocytes (Fig. 2D; Lck-Cre cKO a, CD4-Cre cKO). In marked contrast, we did not observe any histological differences in Lck-Cre cKO mice with normal sized thymi and spleen (Lck-Cre cKO a) or in the thymi of CD4-Cre cKO mice compared to Ctr individuals (Fig. 2D).

### LRH-1 is critical for the transition from DN2/DN3 to DN4 during thymocyte development

Since the 4-HT-inducible and Lck-Cre-mediated deletion of LRH-1 suggests a key role of LRH-1 in early thymocyte development, its impact on individual thymocyte subsets in the Lck-Cre cKO mice was further analyzed by multicolor flow cytometry. Despite the two different phenotypes in the Lck-Cre cKO mice regarding thymus and spleen size, the frequency and absolute numbers of DN thymocytes were significantly altered in all Lck-Cre cKO mice. Moreover, an increased basal cell death of thymocytes was observed (Fig. 3A, Suppl. Fig. 2A-D). The frequency of DN2 and DN3 thymocytes in Lck-Cre cKO mice was increased, whereas DN4 cells were decreased. The frequency of DP and CD4 SP cells remained unchanged, while CD8 SP thymocytes were significantly altered (Fig. 3A). However, the total number of cells in all thymocyte subsets was significantly reduced (Suppl. Fig. 2D). The frequency of GFP^+^ and thus LRH-1-deleted cells was greatly reduced in DN1, DN4, DP, CD4 SP and CD8 SP cells (Fig. 3B), whereas dtTomato^+^ cells were significantly increased in most thymocyte subsets of Lck-Cre cKO mice compared to Ctr mice (Fig. 3B, Suppl. Fig. 2E). Consequently, most of the remaining thymocytes in Lck-Cre cKO mice have escaped Cre-mediated LRH-1 deletion and significantly reduced *Lck* expression (Suppl. Fig. 3A-C). In agreement with this finding, almost no GFP^+^ cells were observed in thymic and splenic tissue sections and in the isolated cells (Suppl. Fig. 2F-G, Suppl. Fig. 3D). Furthermore, splenic T cells of Lck-Cre cKO mice showed no alterations in either *Nr5a2* or *Lck* expression but significantly diminished *Cre* expression (Suppl. Fig. 3A-C). The percentage and the total cell number of thymocytes within the subsets in CD4-Cre cKO mice was significantly reduced for CD4 SP and CD8 SP thymocytes, whereas DN and DP populations remained unaffected (Suppl. Fig. 4A-B). In addition, the total number of mature T cells in the spleen of CD4-Cre cKO mice was also reduced (Suppl. Fig. 4C-D) [8].

**Figure 3:**
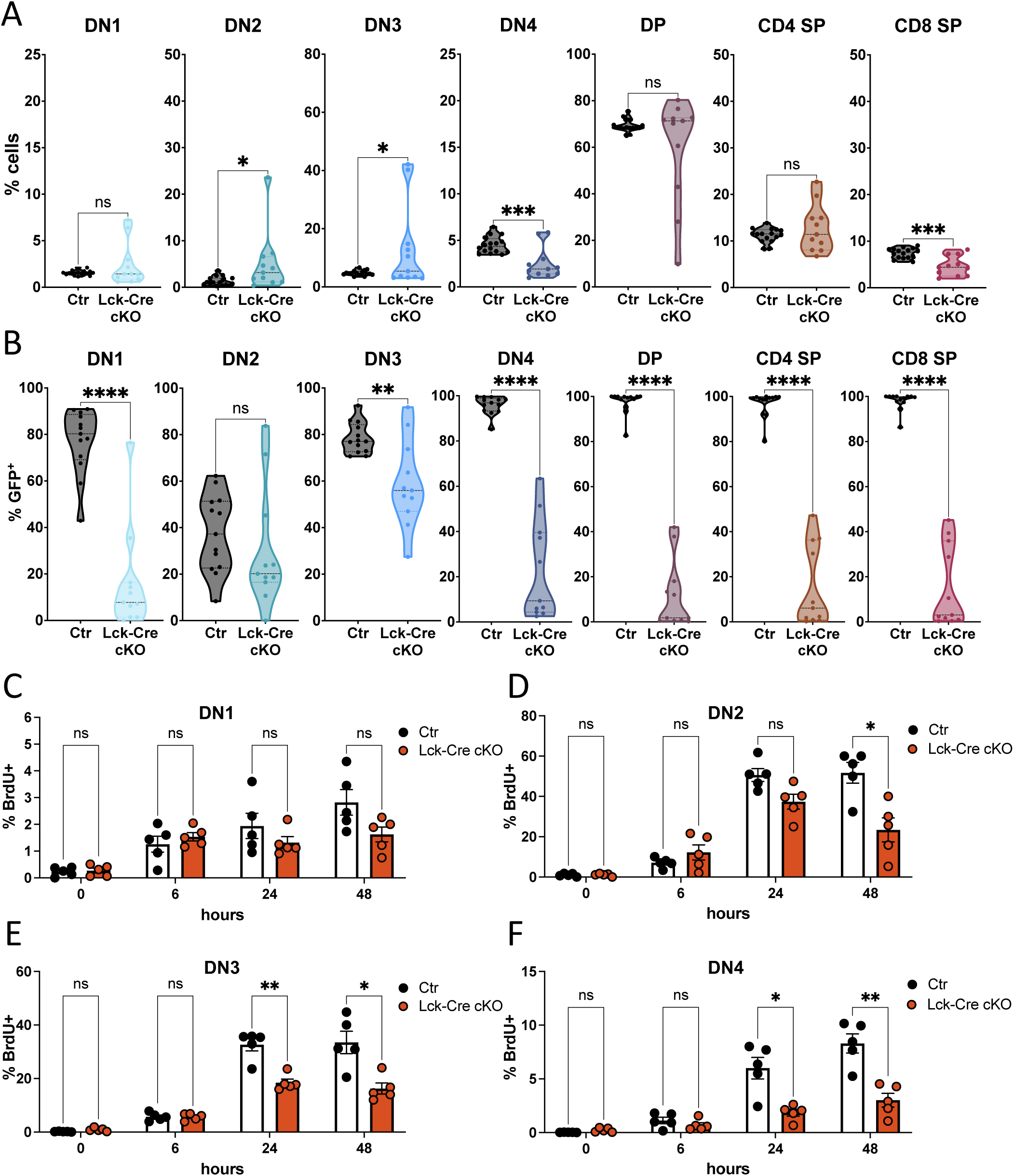
LRH-1 is critical for the transition from DN2/DN3 to DN4 during thymocyte development. High-dimensional flow cytometry analysis of frequency (A) and GFP-positive (B) live thymocyte stages from Ctr and Lck-Cre cKO mice. Thymocyte stages were defined by the expression of CD25, CD44, CD4 and CD8. Each point in the violin plots represents an individual mouse (n = 16 Ctr, n = 11 Lck-Cre cKO). Flow cytometry analysis of proliferating DN1 (C), DN2 (D), DN3 (E) and DN4 (F) thymocytes from Ctr and Lck-Cre cKO mice co-cultured with OP-DL1 cells and treated with IL-7 (1 ng/ml). BrdU-positive live DN1-4 thymocytes at indicated time points are shown. Bars show mean ± SEM, dots represent individual mice (n = 5 per group). Statistical significance was determined by unpaired Student’s t-test (A, B) and multiple unpaired Student’s t-test with Holm-Šídák multiple comparison test (C-F), ns: not significant, * p<0.05, ** p<0.005, *** p<0.001, **** p<0.0001. DN = double negative, DP = double positive, SP = single positive.

The inducible or Lck-Cre-mediated deletion revealed that LRH-1 is critical for early T cell development. Since LRH-1 is known to be a regulator of cell proliferation and two major proliferation waves occur during early murine T cell development, the proliferation in control and Lck-Cre cKO thymocytes was analyzed using BrdU incorporation [6, 8, 9, 15]. The first proliferation wave occurs prior to TCRβ rearrangement in DN1, DN2 and partially in DN3 thymocytes, followed by the second proliferation wave after β-selection in the DN3 to DN4 transition [6, 15]. DN2, DN3 and DN4 thymocytes from Lck-Cre cKO mice showed reduced proliferation at 24 h and/or 48 h after isolation in *ex vivo* co-cultures with OP9-DL1 cells (Fig. 3D-F). In contrast, no changes in BrdU incorporation were observed in DN1, DP, CD4 SP and CD8 SP subsets (Fig. 3C, Suppl. Fig. 3F). Next to reduced proliferation, the basal mitochondrial ATP production of the global population was significantly reduced in Lck-Cre cKO thymocytes (Suppl. Fig. 3E). Thus, although overall *Nr5a2* expression is low in the different DN thymocyte stages, LRH-1 appears to be critical in regulating mitochondrial ATP production and proliferation during early T cell development.

### Reduced frequency of mature CD4^+^ T cells in peripheral lymphoid tissues of Lck-Cre cKO mice

We have previously reported that CD4-Cre-mediated deletion of LRH-1 at the DP thymocyte stage results in a marked reduction of peripheral T cell numbers (Suppl. Fig. 4C-D) and failure of CD4^+^ and CD8^+^ T cell effector functions [8]. Deletion of LRH-1 in the early DN thymocyte stages also resulted in reduced total peripheral T cell numbers, and reduced CD4^+^ T cell frequencies in the spleen and mesenteric lymph nodes (mLN) (Fig. 4A-B, Suppl. Fig. 5A-B). In contrast, peripheral T cells isolated from Peyer’s patches and the intestinal epithelial layer (intraepithelial lymphocytes, IEL), as well as other T cell subsets, such as CD8^+^, regulatory T cells (T_reg_) and γδ T cells from spleen, mLN, Peyer’s patches and the intestinal epithelial layer of Lck-Cre LRH-1 cKO mice did not differ from those of Ctr mice (Fig. 4A-B, Suppl. Fig. 6A-F). Similar to the DN4, DP, CD4 SP and CD8 SP thymocytes, the majority of the remaining peripheral T cells escaped from Lck-Cre-mediated LRH-1 deletion, as indicated by the low numbers of GFP^+^ cells compared to those in Ctr mice (Fig. 4 C-D). Furthermore, CD4^+^ but not CD8^+^ LRH-1 deletion escapers homing to the spleen showed increased basal cell death, but no increased susceptibility to apoptosis induction by etoposide or TCR activation (Suppl. Fig. 5C-E).

**Figure 4:**
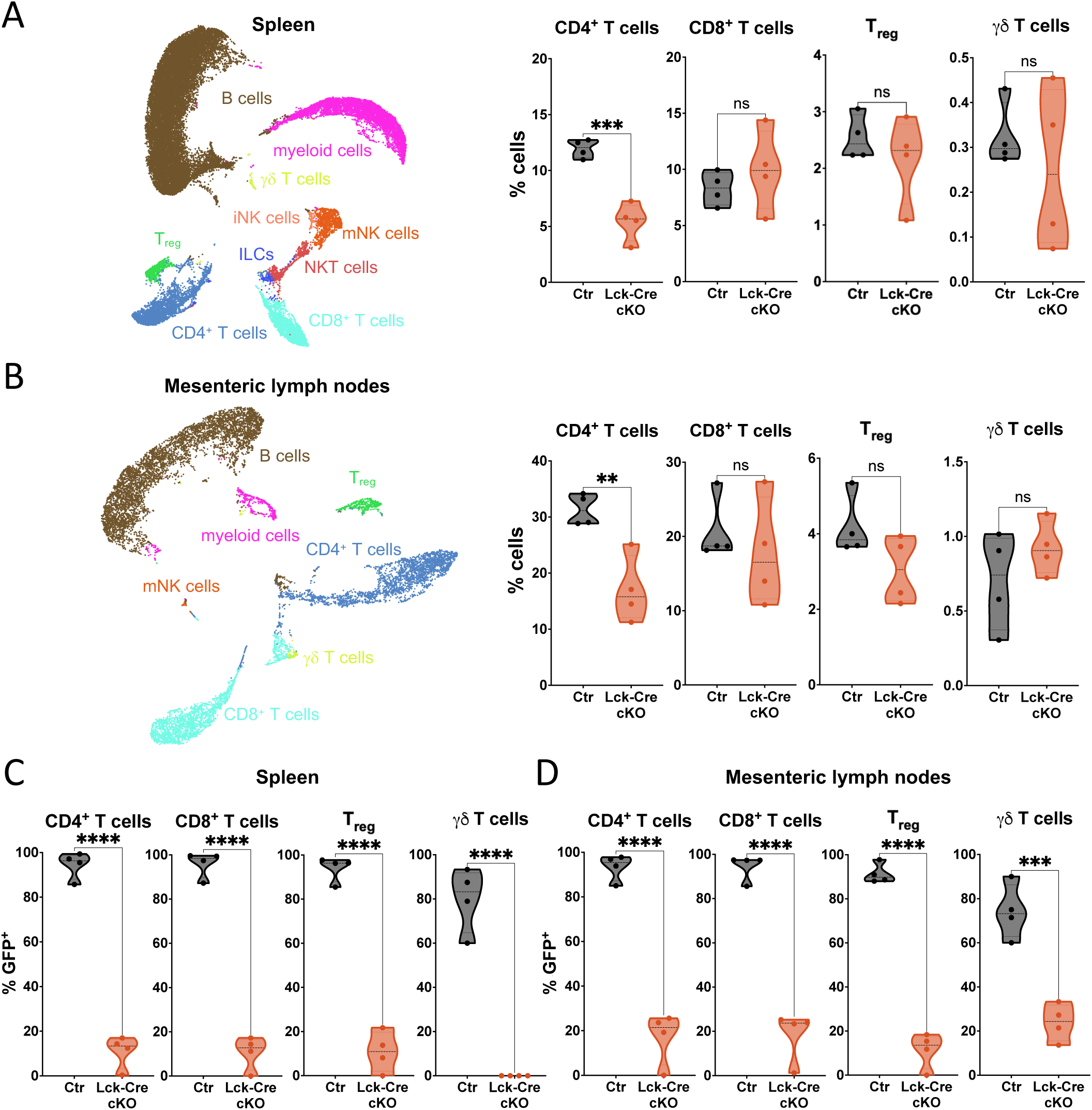
Reduced frequency of mature CD4^+^ T cells in peripheral lymphoid tissues of Lck-Cre cKO mice. UMAP clusters and frequency of high-dimensional flow cytometry analysis of pre-gated CD45-positive live cells isolated from spleen (A) and mesenteric lymph nodes (B) of Ctr and Lck-Cre cKO mice. GFP-positive live T cells isolated from spleen (C) and mesenteric lymph nodes (D). Each point in the violin plot represents an individual mouse (n = 4 per group). Statistical significance was determined by unpaired Student’s t-test, ns: not significant, ** p<0.005, *** p<0.001, **** p<0.0001. T_reg_ = regulatory T cells, γδ T cells = gamma-delta T cells, iNK cells = immature natural killer cells, mNK cells = mature natural killer cells, NKT cells = natural killer T cell- like cells, ILCs = innate lymphoid cells.

### Impaired activation marker expression and cytokine secretion in LRH-1 deletion escape T cells

To investigate whether T cells escaping Lck-Cre-mediated LRH-1 deletion mature into fully functional peripheral T cells, early and late activation markers, proliferation and pro-inflammatory cytokine secretion in response to TCR stimulation were analyzed. CD8^+^ and CD4^+^ T cells showed delayed and strongly reduced expression of the activation markers CD69 and CD25 after TCR stimulation (Fig. 5A-B). However, the proliferation of splenic CD4^+^ and CD8^+^ T cells did not differ from that of the Ctr T cells (Fig. 5C-D), whereas basal mitochondrial ATP production under steady-state conditions and after T cell activation was reduced (Suppl. Fig. 5F). Consistent with our previous study, splenic T cells from Lck-Cre cKO mice showed a significantly increased release of the T_H_1/T_C_1 cytokines IL-2, TNF and IFN-γ, as well as IL-6 after TCR stimulation (Fig. 5E-H), but no differences in the secretion of the T_H_2 cytokine IL-4 (Suppl. Fig. 5G) [8]. Thus, peripheral T cells from Lck-Cre cKO mice do not appear to be anergic, but show reduced expression of surface activation markers and a shift toward T_H_1/T_C_1 T cells upon TCR stimulation.

**Figure 5:**
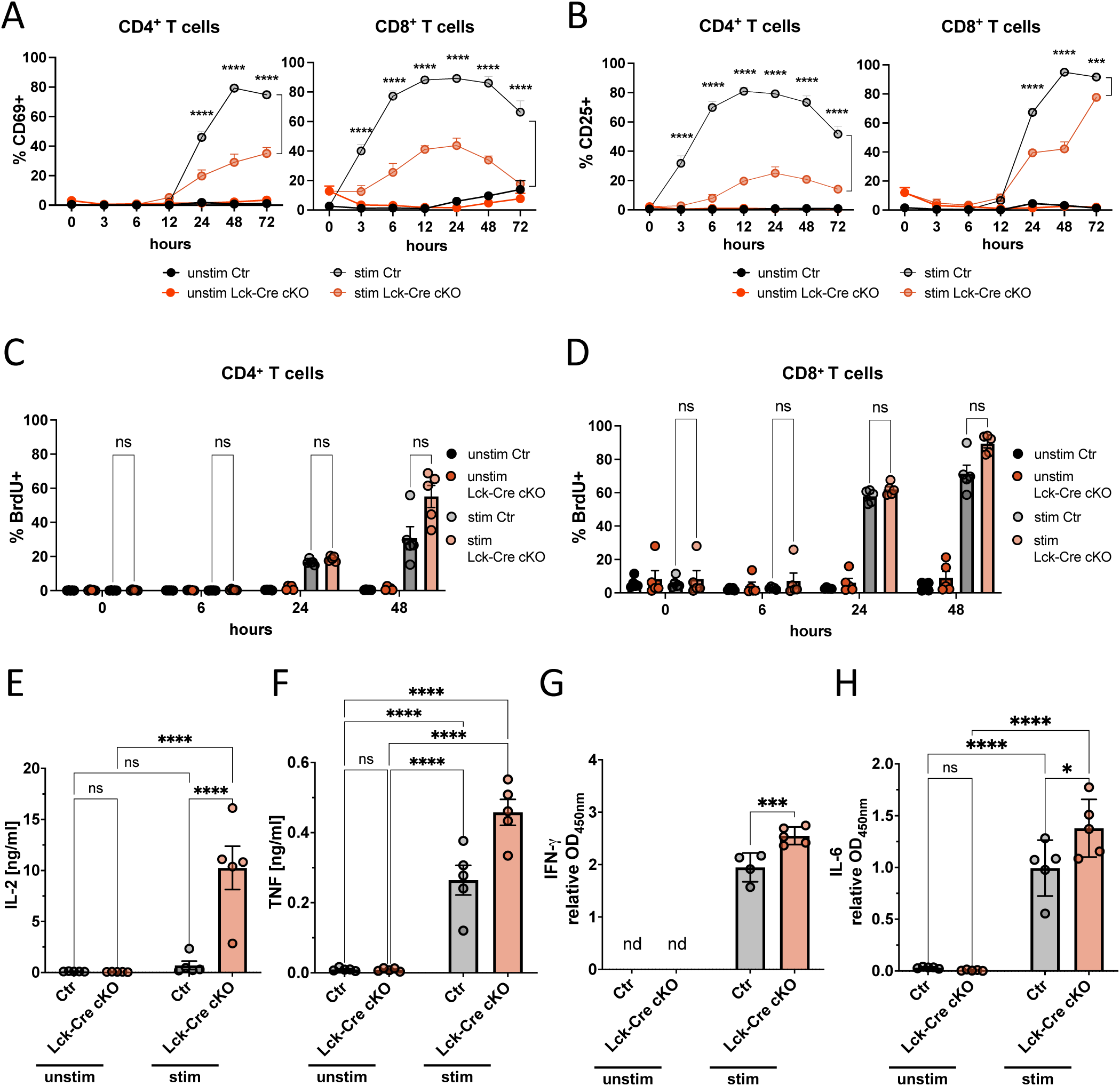
Impaired activation marker expression and cytokine secretion in LRH-1 deletion escape T cells. Flow cytometry analysis of CD69-positive (A) and CD25-positive (B) CD4^+^ and CD8^+^ T cells, after stimulation or unstimulated, at indicated time points. Points represent mean ± SEM (n = 3-4 mice). Flow cytometry analysis of BrdU-positive live stimulated or unstimulated CD4^+^ T cells (C) and CD8^+^ T cells (D) from the spleen of Ctr and Lck-Cre cKO mice. Bars show mean ± SEM, dots represent individual mice (n = 5 per group). Secretion of IL-2 (E), TNF (F), IFN-γ (G) and IL-6 (H) in unstimulated and stimulated splenocytes. Points represent mean ± SEM (n = 5 mice). (A-H) T cells were stimulated with plate bound anti-CD3ε (3 µg/ml) and anti-CD28 (1 µg/ml) or unstimulated for 48 h. (E-H) Primed cells were restimulated after 42 h with PMA (1 ng/ml) and ionomycin (200 ng/ml) for 6 h. Statistical significance was determined by multiple unpaired Student’s t-test with Holm-Šídák multiple comparison test for stim Ctr and stim Lck-Cre cKO (A-D) and ordinary two-way ANOVA with Turkey multiple comparison test (E-H), nd = not detected, ns: not significant, * p<0.05, *** p<0.001, **** p<0.0001. Unstim = unstimulated, stim = stimulated.

### CD4^+^ T cells from LRH-1 Lck-Cre cKO mice fail to induce effector T cell-mediated colitis and impair regulatory T cell-mediated protection

Although we have identified significant changes in early T cell proliferation, mature T cell surface marker expression and T_H_1/T_C_1 cytokine secretion after T cell stimulation in LRH-1 Lck-Cre cKO mice, the overall functionality of T cells from the Lck-Cre cKO mice, especially CD4^+^ T cells, remained unclear. To address this question, naïve effector T cells (CD4^+^ CD45RB^hi^; T_Eff_) were transferred into Rag2^-/-^ immunodeficient mice to establish T_Eff_-mediated experimental colitis. Moreover, the functionality of regulatory T cells (CD4^+^ CD45RB^lo^; T_reg_) was examined in a T_reg_-mediated protection from T_Eff_-induced colitis [16, 17]. The transfer of T_Eff_ from the Lck-Cre Ctr mice led to infiltration of the colon, colonic inflammation and consequently a steady decrease in Rag2^-/-^ recipient mice’s body weight. T_Eff_ from the Lck-Cre cKO mice failed to promote inflammation and no body weight loss was observed (Fig. 6A, Suppl. Fig.7E top panel). At the same time, T_Eff_-injected Rag2^-/-^ mice co-transferred with T_reg_ from Ctr and wildtype B6 mice were protected from colitis. In contrast, T_reg_ from the Lck-Cre-cKO mice failed to prevent colitis-associated body weight loss and led to cell infiltration in the colon accompanied with colonic inflammation (Fig. 6A, Suppl. Fig. 7E top panel). Rag2^-/-^ recipient mice transferred with Lck-Cre cKO T_Eff_ showed reduced CD4^+^ and IL17A^+^ T cell numbers in the colonic LP, mLN and spleen, as well as IFN-γ^+^ and IL17A^+^IFN-γ^+^ T cells in the LP, compared to the control T_Eff_ transfer group. In the T_reg_ co-transfer model, cell numbers in all T cell populations were increased in the Lck-Cre cKO transfer group compared to the controls and reached levels similar to the Ctr T_Eff_ transfer group. Additionally, total T_reg_ numbers in the Lck-Cre cKO T_reg_ co-transfer group remained lower than in both co-transfer controls (Fig. 6B-C, Suppl. Fig. 7A, D). In agreement with previous experiments, GFP expression of transferred T cells from Lck-cKO mice was reduced in the LP and mLN of Rag2^-/-^ mice compared to T cells transferred from Ctr mice, while T cells transferred from B6 mice exhibited no detectable GFP expression (Suppl. Fig.7 B-C). Additionally, serum levels of the pro-inflammatory IFN-γ and TNF cytokines, as well as *Ifng* and *Tnf* mRNA expression in the colon were decreased in Rag2^-/-^ mice receiving the Lck-Cre cKO T_Eff_ compared to those receiving Ctr T_Eff_. In contrast, these inflammation parameters were all increased in the co-transfer mice receiving the Lck-Cre cKO T_reg_ (Fig. 6 D-E). These findings indicate that CD4^+^ T_Eff_ and T_reg_ from Lck-Cre cKO escaping Cre-mediated LRH-1 deletion are functionally impaired in T cell-mediated intestinal inflammation.

**Figure 6:**
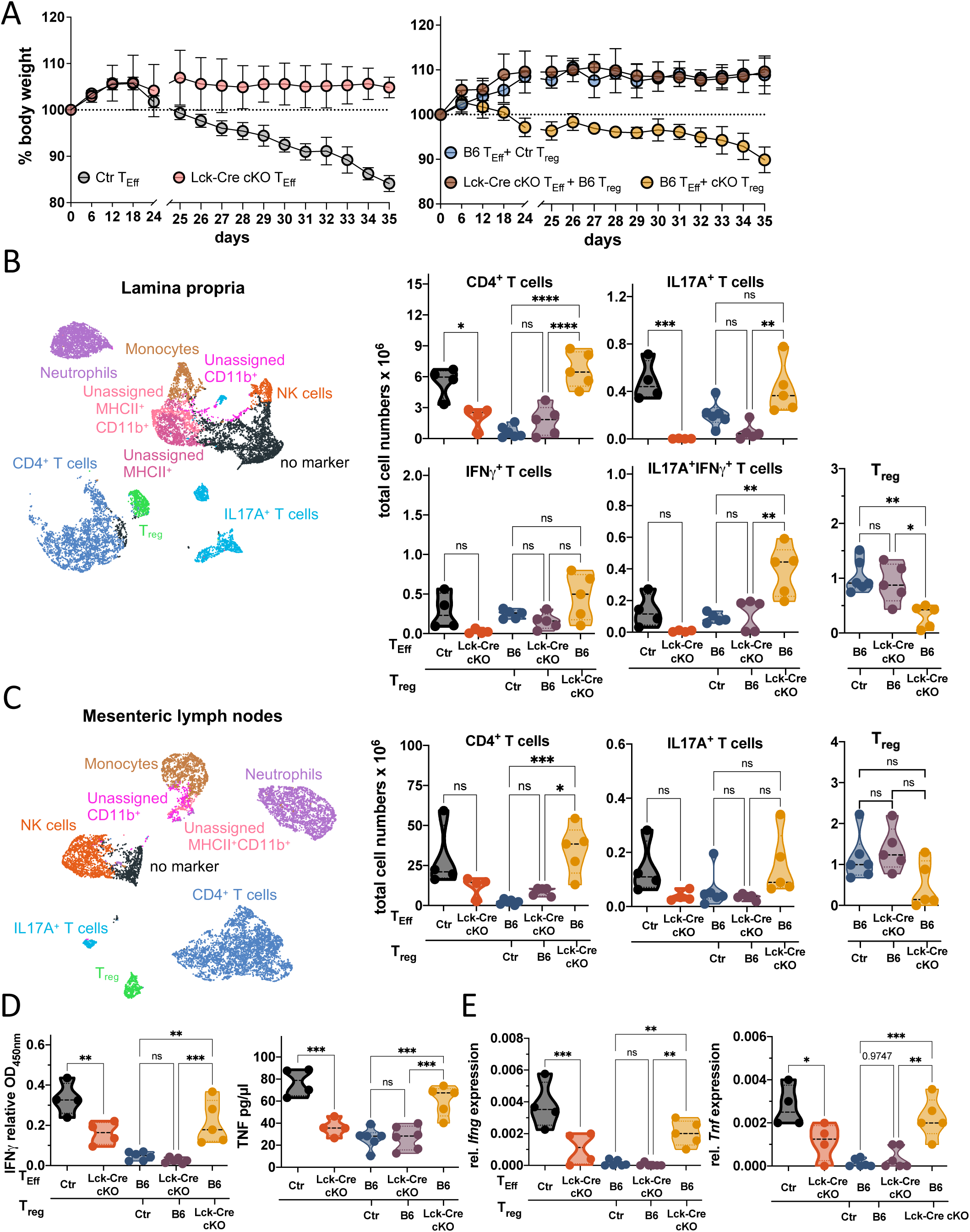
CD4^+^ T cells from LRH-1 Lck-Cre cKO mice fail to induce effector T cell-mediated colitis and show impaired regulatory T cell-mediated protection. (A) Body weight changes over time in Rag2^-/-^ mice after transfer of T_Eff_ and co-transfer of T_Eff_ and T_reg_ from Lck-Cre cKO, Lck-Cre x mTmG control mice (Ctr) or wildtype mice (B6). UMAP clusters of immune cells and total T cell numbers x 10^6^ from colonic lamina propria (B) and mesenteric lymph nodes (C). Data were pre-gated to live CD45^+^ cells. TNF and IFN-γ secretion in the serum (D) and relative RT-qPCR of *Ifng* and *Tnf* expression in the colon (E) of Rag2^-/-^ mice after T_Eff_ transfer or T_Eff_ and T_reg_ co-transfer. Gene expression was normalized to *Actb*. Each point in the violin plots represents an individual mouse (n = 4 T_Eff_ transfer of Ctr and Lck-Cre cKO, n = 5-6 T_Eff_ and co-transfer of Ctr, B6 and Lck-Cre cKO T_reg_. Statistical significance was determined by one-way ordinary ANOVA with Dunnett’s multiple comparison test, ns: not significant, * p<0.05, ** p<0.005, *** p<0.001, **** p<0.0001.

## Discussion

The pleiotropic functions of LRH-1 regulating development, stemness, proliferation and metabolism have been extensively studied in various tissues of endodermal origin, where LRH-1 is abundantly expressed. In contrast, only a limited number of studies have focused so far on the roles of LRH-1 in the hematopoietic system. This is particularly true for lymphocytes and the T cell lineage. At least in part, this limited interest is related to the relatively low LRH-1 expression in these cells. Nevertheless, even though LRH-1 expression is modest compared to that in the liver, pancreas or intestine, it appears to regulate critical processes in T cells, such as proliferation, cytokine secretion, and cell death [8, 18, 19]. Of note, while hepatocyte-specific deletion results in the development of a metabolic syndrome, the liver still has normal architecture, cellular composition and size [20, 21]. The deletion of LRH-1 in the T cell lineage has far more severe consequences for the development or immature and mature T cells. In a previous study, we have shown that CD4-Cre-mediated deletion of the *Nr5a2* gene had little impact on thymocyte frequency, yet resulted in strongly reduced numbers of mature T cells and severely impaired effector functions [8], clearly highlighting the crucial role of LRH-1 in regulating mature T cell response. Although immature T cells exhibit similar LRH-1 expression levels as mature T cells [22] (Fig. 1A), the potential relevance of LRH-1 in thymic development had not previously been investigated. To address the relative importance of LRH-1 at different thymocyte stages, an inducible LRH-1 deletion system was established, in which thymocytes were co-cultured with OP9-DL1 cells to mimic the thymic niche and enable T cell development *ex vivo* [11, 23]. Because Cre activation and/or treatment with high concentrations of the functional metabolite of tamoxifen (4-hydroxytamoxifen; 4-HT) are known to be toxic to immune cells, we used a concentration of 4-HT that resulted in only partial LRH-1 deletion, but no adverse effects on the control thymocytes [24–26] (Suppl. Fig. 1E). Our study revealed that *Nr5a2* is expressed at elevated levels in DP and CD4 SP thymocytes. However, deletion of LRH-1 did not significantly affect their relative frequency. This finding is consistent with our previous study, in which CD4-Cre-mediated LRH-1 deletion showed no effect on the distribution of DP, and CD4 or CD8 SP thymocytes [8]. In contrast, DN thymocytes express lower levels of LRH-1 than DP and SP thymocytes, yet LRH-1 appears to be more crucial for their further development. Thus, we found that the proportion of DN3 and DN4 cells after 4-HT-induced LRH-1 deletion decreased significantly, indicating a potential role for LRH-1 in early thymocyte development. This idea was further confirmed by Lck-Cre-mediated LRH-1 deletion, which revealed a significant increase in DN2 and DN3 cell abundance, but a decrease in DN4 thymocyte numbers. The contrasting relative distribution of the DN populations in the two LRH-1 knockout models may be attributed to differences in the experimental setup and knockout efficiency. In the inducible LRH-1 deletion system, the frequencies of thymocytes were analyzed *ex vivo* 72 hours after isolation without replenishment by bone marrow–derived precursors. In contrast, the frequency of thymocyte subsets in Lck-Cre cKO mice was directly assessed after isolation. Additionally, Cre activation, and thus LRH-1 deletion, was found to be only partially effective in the *ex vivo* inducible knockout system, albeit most effective in DN2, DN3, and DN4 thymocyte stages. In the *in vivo* Lck-Cre-driven LRH-1 knockout mouse model, Cre activation was most efficient in DN2 and DN3 thymocyte stages. Low levels of GFP expression, monitoring Cre activation in all thymocyte subsets beyond the DN3 stage, as well as in peripheral T cells, and equal *Nr5a2* expression in thymocytes and splenic T cells from the Lck-Cre LRH-1 cKO and Ctr mice, indicate unsuccessful Lck-Cre-mediated LRH-1 deletion in our cKO mouse model. Expression of Cre recombinase driven by the proximal Lck promoter is commonly used for gene deletion at the early DN thymocyte stage, and is only insufficient in the Lck-Cre LRH-1 cKO but not in the Lck-Cre Ctr mice [27].

Instead, our data rather suggest that loss of LRH-1 at these early thymocyte stages is detrimental for further thymic development and progression to mature peripheral T cells. Thus, the majority of immature and mature T cells appears to shut down Cre expression in order to prevent LRH-1 deletion. A number of studies have indicated that silencing of the Cre expression can be attributed to epigenetic modifications, particularly DNA methylation in the promoter region [28–30]. Other potential causes of shutting down Cre expression, such as position effect variegation [29, 31] or limited accessibility of the floxed genes are unlikely, as the Ctr mice do not show impaired Cre activation. In agreement with this notion is the fact that most thymocytes beyond DN3 stage and peripheral T cells remained dtTomato^+^ and thymocytes had low levels of *Lck* expression. Although not formally proven, this suggest that silencing of the Lck proximal promoter, driving Cre transgene expression as well as endogenous Lck expression in thymocytes, allows T cells to prevent the unfavorable deletion of LRH-1 and associated consequences in mice. The *Lck* transcript is not altered in mature T cells, which suggest that the Lck distal promoter is not silenced in peripheral T cells [32, 33].

The question remains which LRH-1-regulated target genes and processes are essential for thymic T cell development. Previous studies have highlighted the critical role of LRH-1 in regulating the cell cycle and proliferation via the transcriptional control of cyclin E1, cyclin D1, and c-Myc. [8, 34–36]. Accordingly, we reported previously that CD4-Cre-mediated LRH-1 deletion or pharmacological inhibition of LRH-1 decreased activation-induced proliferation of CD4^+^ and CD8^+^ T cells [8]. Similarly, LRH-1 inhibition or downregulation also inhibited proliferation of human leukemic T cells [9]. Conversely, Cobo-Vuilleumier et al. observed diminished CD4^+^ and CD8^+^ T cell proliferation in peripheral blood mononuclear cells of type 1 diabetes patients upon LRH-1 activation, rather than inhibition [8, 19]. In our present study, LRH-1 deletion in DN thymocytes resulted in fewer BrdU^+^ DN2, DN3, and DN4 cells, indicating reduced DNA synthesis, though no differences were observed at later stages. The development of immature T cells is characterized by two major waves of proliferation at the DN stage, and clonal expansion at the DP stage. The first wave occurs before TCRβ selection in DN1 to early DN3 cells. The second wave occurs after TCRβ gene rearrangement in late DN3 to DN4 thymocytes and is followed by pre-TCR expression [5, 15, 37–39]. c-Myc is important not only for the proliferation of mature T cells, but also in mediating pre-TCR-induced proliferation of DN thymocytes [8, 34, 35]. The increased proportion of DN3 thymocytes, the high level of Cre activation similar to the control and the significant reduction of DN4 GFP^+^ cells in Lck-Cre-mediated LRH-1 cKO support the hypothesis that LRH-1 deletion results in proliferation defects in pre-TCR-expressing DN thymocytes. This effect is likely mediated by impaired expression of proliferation-regulating genes, such as c-Myc, thereby preventing the transition from the DN3 to DN4 stage.

T cell proliferation initiated by TCR activation is also closely linked to a rapid metabolic switch from oxidative phosphorylation to glycolysis, which also requires c-Myc signaling [34, 35, 40–43]. A similar metabolic switch occurs in the development of DN thymocytes, where DN3 cells primarily rely on oxidative phosphorylation, while DN4 cells mostly depend on glycolysis [44, 45]. LRH-1 not only regulates c-Myc expression but also modulates glutaminase 2 (GLS2) expression and activity. GLS2 is an enzyme important for glutamine-dependent α-ketoglutarate production and associated mitochondrial ATP production [46]. In our present study, we show that thymocytes and peripheral T cells from Lck-Cre LRH-1 cKO mice have reduced basal, mitochondrial-dependent ATP production, and no increase in ATP production could be observed in CD4^+^ and CD8^+^ T cells upon TCR stimulation. Other key regulators of early T cell development and proliferation include c-Kit, IL-7, and Notch signaling [47–51]. DN1 and DN2 thymocytes require IL-7 and Notch signaling for proliferation, while DN3 thymocytes rely solely on Notch activation [48]. Notch signaling leads to the induction of TCF-1, one of the first T cell-specific genes during T cell development. TCF-1 is a transcription factor that belongs to the WNT-β-catenin signaling cascade, a pathway that is critical for the development of DN thymocytes [52–54]. Notably, LRH-1 was found to interact also with the WNT-β-catenin signaling cascade by heterodimerizing with β-catenin, and transcriptionally regulating cyclin E1 and cyclin D1 expression [36]. Further evidence of a potential link between LRH-1 and the WNT-β-catenin pathway comes from a publication by Xu et al., who investigated Lck-Cre-mediated β-catenin deletion in mice [55]. Interestingly, both the Lck-Cre β-catenin and Lck-Cre LRH-1 cKO mice show an impaired transition from the DN3 to the DN4 stage, and reduced peripheral T cell numbers, suggesting common regulatory circuits. Taken together, LRH-1 deletion results in impaired DN thymocytes development, failing to progress beyond the DN3 stage. Likely, this defect is related to the important role of LRH-1 in regulating cell cycle progression and proliferation, but may include, however, also pro-survival functions of LRH-1, as observed in other cellular systems [8, 34–36, 56–59].

Our LRH-1 deletion experiments reveal that LRH-1 expression exerts substantial selection pressure during thymocyte development, resulting in preferential selection of thymocytes that escape Cre expression. Interestingly, while in some mice Lck-Cre-mediated LRH-1 deletion resulted in a reduced thymus and spleen size, in other mice thymus and spleen size, and histology appeared normal, although they were primarily populated by dtTomato^+^ Cre escapers. Since these cells have an intact LRH-1 gene, one would expect normal TCR-induced proliferation and effector functions. Indeed, peripheral CD4^+^ and CD8^+^ T cells do not exhibit proliferation defects, however, a skewed activation and cytokine profile. Upon TCR activation, LRH-1-deficient T cells release massive amounts of IFN-γ, IL-2, and TNF, while showing reduced expression of early T cell activation surface markers. Interestingly, similar results were observed in CD4-Cre-mediated LRH-1 cKO T cells, which showed strongly increased IFN-γ and IL-2 secretion [8]. This suggests a crucial role for LRH-1 in T helper cell differentiation, respectively T_H_1/T_H_2 cytokine balance. In agreement with this notion is the observation that LRH-1 activation in human T cells from patients with type 1 diabetes results in decreased IFN-γ production, and a shift toward anti-inflammatory T_reg_ and a T_H_2-like phenotypes [19]. Also, TCF-1 deficiency in mature CD4^+^ T cells promotes increased IFN-γ production, as TCF-1 and β-catenin regulate T_H_2 lineage differentiation by binding to the promoter region of GATA-3 [60]. Although peripheral T cells from Lck-Cre LRH-1 cKO mice have normal *Nr5a2* and *Lck* expression levels as they escaped LRH-1 deletion, they have a marked reduction in *Lck* expression during T cell development. Lck deficiency in T cells is known to impair T_H_2 cell differentiation, promote IFN-γ production in T_H_2-polarized cells, and decrease IL-4 production [61, 62]. Lck expression is regulated by a proximal and a distal promoter. The deletion of the distal Lck promoter reduces CD4^+^ T cell proliferation upon anti-CD3 stimulation, resembling the phenotype of complete Lck knockout T cells, whereas deletion of the proximal Lck promoter does not affect peripheral T cell proliferation [33]. Therefore, the impaired and delayed expression of T cell activation markers, together with the shift toward T_H_1/T_C_1 cytokine secretion upon TCR stimulation in Lck-Cre LRH-1 cKO T cells, is likely due to defective maturation processes during thymocyte development. The absence of activation-induced T cell proliferation defects and the unchanged *Lck* expression in peripheral T cells support the hypothesis, that these observations are not caused by LRH-1 or Lck deficiency in mature T cells. The study of single Lck promoter deletion further demonstrated that both promoters are essential for normal thymocyte development, and consequently functional mature T cells [33]. Consistent with these findings, both the T_Eff_-mediated colitis model and the T_reg_-mediated protection against T_Eff_-mediated colitis demonstrate that CD4^+^ Lck-Cre cKO T cells exhibit compromised effector and regulatory functions during T cell-mediated immune responses.

Taken together, our results demonstrate the important role of LRH-1 in thymocyte maturation, especially in early DN thymocytes. We propose that the absence of LRH-1 during the initial waves of thymocyte proliferation impairs the ability of DN thymocytes to expand and differentiate beyond the DN3 stage. Likely, this is due to impaired LRH-1-regulated transcriptional regulation of thymocyte metabolism, cell cycle, and proliferation. Possibly, this process may also involve heterodimerization and cooperation of LRH-1 with other transcription factors involved in thymocyte differentiation, such as β-catenin [36, 63]. The pressure to maintain LRH-1 expression during thymic selection up to the DN3 stage appears to be so high, that most thymocytes at the DN4 stage or later are Cre escapers to maintain LRH-1 expression. Thus, the Lck-Cre-mediated LRH-1 cKO mouse model does not allow to directly compare the relative role of LRH-1 in thymocytes versus mature T cells. Yet, this study on Lck-Cre-mediated LRH-1 deletion and our past study on CD4-Cre-mediated LRH-1 deletion strongly supports the notion that LRH-1 is critical for both, immature and mature T cell development and function [8]. This strong impact of LRH-1 deletion in the T cell lineage is rather surprising given that T cells express very low levels of LRH-1 compared to the liver or intestine. At the same time, this low but critical expression of LRH-1 in the T cell lineage may also offer an interesting therapeutic window, in which T cell-mediated pathologies may be targeted pharmacologically, while avoiding deregulation of various other LRH-1-regulated homeostatic processes in different organs. Along these lines, it is interesting to note that a pharmacological inhibitor of LRH-1 protected from T cell-mediated hepatitis yet had only minimal impact on normal liver integrity [8, 18, 19].

## Material and Methods

### Mice

All animal experiments and isolation of primary cells were conducted in accordance with the German Animal Welfare Regulations and approved by the Ethics Committee of the Regierungspräsidium Freiburg i.B.. All mice were housed in individually ventilated cages under specified pathogen-free conditions with a 12 h light-dark cycle in the central animal facility of the University of Konstanz. Mice had access to food and water ad libitum. For all experiments, mice were used with a similar distribution of females and males aged between 7 and 16 weeks. C57BL/6 mice (B6) and mice generated with the floxed Liver receptor homolog-1 (LRH-1) gene (LRH-1^L2/L2^), CD4-Cre, lymphocyte protein tyrosine kinase (Lck)-Cre, Rosa26Cre-ERT2, Rag2^-/-^ and the two-color fluorescence reporter construct containing floxed membrane-targeted tandem dimer Tomato (dtTomato) followed by membrane-targeted GFP (mTmG) have been reported previously [12, 14, 64–66].

### OP9-DL1 Thymocyte Co-culture System

The murine bone marrow-derived stromal cell line OP9-DL1 [23] was cultured in Iscove’s modified Dulbecco’s medium containing 5 % fetal bovine serum (FBS), 20 µg/ml gentamycin, 2 mM L-glutamine and 50 µM 2-mercaptoethanol, 1x MEM non-essential amino acids, 0.03 % pepton and 1x insulin transferrin selenium. The protocol for the co-culture of thymocytes on OP9-DL1 cells was adapted from Holmes and Zúñiga-Pflücker [11]. Briefly, OP9-DL1 cells were seeded at a density of 4.5 x 10^4^ cells/cm^2^ one day prior to thymocyte isolation. Subsequently, 4 x 10^5^ thymocytes/ml in Roswell Park Memorial Institute (RPMI) 1640 medium containing 5% FBS, 20 µg/ml gentamycin, 2 mM L-glutamine, 50 µM 2-mercaptoethanol and 1 ng/ml murine interleukin (mIL-) 7 were added for subsequent culture up to 72 h.

### Isolation of Thymocytes, Splenocytes and Immune Cells from Mesenteric Lymph Nodes and Peyer’s Patches

Immune cells from thymus, spleen, mesenteric lymph nodes (mLN) and Peyer’s patches were released in phosphate-buffered saline (PBS) by crushing the tissues between frosted glass slides. After erythrocyte lysis in PBS, 150 mM NH4Cl, 10 mM KHCO_3_ and 100 µM Na_2_ ethylenediaminetetraacetic acid (EDTA) for splenocyte and thymocyte preparations, immune cells were counted and cultured in RPMI 1640 containing 5% FBS, 20 µg/ml gentamicin, 2 mM L-glutamine and 50 µM 2-mercaptoethanol, or directly transferred to Flow Cytometry (FC) Buffer containing PBS, 2.5% bovine serum albumin (BSA), 2 mM EDTA and 0,01% sodium azide.

### Isolation of Intraepithelial Lymphocytes

Intraepithelial lymphocytes (IEL) were isolated from the small intestine as previously described [67, 68]. Briefly, epithelial cells and IEL were dissociated in Hanks balanced salt solution (HBSS) and 4-(2-hydroxyethyl)-1-piperazineethanesulfonic acid (HEPES)-bicarbonate buffer (HHB) containing 2% horse serum (HS), 1 mM dithiothreitol (DTT) and 500 µM EDTA, and further separated by Percoll density centrifugation (40%/70%). The interphase containing IELs was harvested, washed in HHB containing 5% HS and resuspended in FC Buffer for flow cytometry analysis.

### Isolation of Lamina Propria Lymphocytes

Lamina propria lymphocytes (LPL) were isolated from the colon as previously described with adaptations [67, 69]. Briefly, epithelial cells from the colon were removed using HHB containing 2% HS, 1 mM DTT and 500 µM EDTA. To release lymphocytes from the lamina propria, remaining colon pieces were enzymatically digested using HHB containing 5% HS, 1.3 mM CaCl_2_, 0.5 mM MgCl_2_, 0.6 mM MgSO_4_, 1 mg/ml collagenase IV and 100 µg/ml DNase I. LPL were separated by Percoll density centrifugation (40%/70%). The interphase containing LPL was harvested, washed in HHB containing 5% HS and resuspended in FC Buffer for flow cytometry analysis.

### Splenic T cell Activation and Cell Death Assays

For T cell activation, isolated cells were stimulated with plate-bound anti-CD3ε (clone: 145-2C11) antibody (3 µg/ml) and soluble anti-CD28 (clone 37.51) antibody (1 µg/ml) to measure activated T cell surface markers by flow cytometry, and additionally restimulated for 6 h with phorbol-12-myristat-13-acetate (PMA) (1 ng/ml) and ionomycin (200 ng/ml) for cytokine detection in the supernatant by ELISA. For basal ATP production rate measurements, CD4^+^ and CD8^+^ T cells were isolated using biotinylated anti-CD4 and anti-CD8 antibodies and magnetic streptavidin beads. For cell death assays, isolated splenocytes were cultured with or without etoposide for 6 h or with plate-bound anti-CD3ε for 24 h, followed by 4′,6-diamidino-2-phenylindole (DAPI) staining and flow cytometry measurement.

### T cell Transfer Model of Colitis

Effector T cell- and regulatory T cell-mediated responses were analyzed in the T cell transfer model of experimental colitis as previously described [16, 17]. Effector T cells (CD4^+^CD45RB^hi^; T_Eff_) and regulatory T cells (CD4^+^CD45RB^lo^; T_reg_) were isolated and sorted from splenocytes of Lck-Cre cKO or control mice (Lck-Cre; Ctr or C57BL/6 wildtype; B6) using the antibodies listed in Reagents and Tools table (RTT). A minimum purity of 98% in post-sorted cells was verified. For the induction of T_Eff_-mediated colitis, 0.5 x 10⁶ T_Eff_ were injected intraperitoneally (i.p.) into Rag2^-/-^ mice. Colitis development was monitored by recording body weight changes. Mice were euthanized after loss of ≤ 20% body weight. To evaluate the functionality of T_reg_, 0.5 x 10⁶ T_Eff_ from control B6 mice were co-injected with 0.2 x 10⁶ T_reg_ from Lck-Cre cKO mice into Rag2^-/-^ recipients. As controls, T_reg_ from Ctr or B6 mice were co-transferred with T_Eff_ from B6 or Lck-Cre cKO mice. These mice were analyzed at the same timepoint as the single T_Eff_ transfer group. In both experiments, immune cells from secondary lymphoid organs and lamina propria (LP) isolated from one-third of the colon were analyzed by flow cytometry. The remaining two-thirds of the colon were divided for histological analysis and RNA extraction, respectively. Serum was collected from Rag2^-/-^ mice by cardiac puncture, followed by centrifugation of whole blood at 2’500 x g for 15 min.

### Flow Cytometry Staining and Fluorescence-activated Cell Sorting

Immune cell suspensions from thymus, spleen, mesenteric lymph nodes, Peyer’s patches, IEL and LPL were stained with different antibody panels listed in RTT. In general, cells were blocked with TruStain FcX™ Plus antibody in FC buffer for 30 min at 4°C. For intracellular antibody staining, cells were first incubated in fixative viability dye (Thermo Fisher Scientific) for 30 min at 4°C in the dark. Cells were then added to the appropriate antibodies in FC buffer and incubated for 45 min at 4°C in the dark. After two washes, cells for intracellular staining were fixed and permeabilized with Transcription Factor Staining Buffer Set (Thermo Fisher Scientific) according to the manufacturer’s protocol and stained with the indicated intracellular antibodies for 45 min at 4°C in the dark. For the bromodeoxyuridine (BrdU) proliferation assay, DNAse I digestion was performed at 37°C in the dark for 1 h prior to intracellular staining. Cells stained with extracellular antibodies only were incubated with DAPI for 10 min at 4°C in the dark. Samples were analyzed on an LSR Fortessa (Becton Dickinson). For fluorescence-activated cell sorting, antibody- and DAPI-stained thymocytes were resuspended in sorting buffer containing PBS, 1 mM EDTA, 25 mM HEPES pH 7, and 1% BSA, filtered through a 30 µm strainer, and sorted on a FACS Aria III sorter (BD). Live thymocytes (DAPI^-^) sorted for double negative 1 (DN1) (CD4^-^ CD8^-^) (CD44^+^ CD25^-^), DN2 (CD44^+^ CD25^+^), DN3 (CD44^lo^ CD25^+^), DN4 (CD44^-^ CD25^-^), double positive (DP) (CD4^+^ CD8^+^), single positive (SP) CD4 (CD4^+^) and SP CD8 (CD8^+^) populations and were subsequently used for RT-qPCR analysis. In addition, for the colitis mouse model experiments, T_Eff_ and T_reg_ were sorted from live (DAPI^-^) splenocytes based on their CD4^+^CD45RB^hi^ and CD4^+^CD45RB^lo^ surface marker expression following the same staining protocol.

### High-Dimensional Flow Cytometry Analysis

Immune cell phenotyping was performed by high-dimensional flow cytometry analysis. Cells were stained with antibody panels listed in RTT, and subsequently analyzed on an LSR Fortessa flow cytometer (BD). FCS files were compensated, cleaned, and pre-gated for single cell, live (DAPI^-^), and CD45^+^ populations using FlowJo (v10.8.1) analysis software. FCS data were exported and analyzed in R using an adapted R script based on previous published studies [70–72]. FCS files were transformed using the inverse hyperbolic arcsinh function and surface expression markers were normalized between values of 0 and 1 with low (1%) and high (99%) percentiles as a limit. Normalized data were applied to Uniform Manifold Approximation and Projection (UMAP) analysis (umap package v0.2.8.0) and unsupervised FlowSOM clustering (FlowSOM package v2.0.0 and ConsensusClusterPlus v1.56.0). The generated clusters were manually merged and annotated based on the pattern of expression markers to different immune cell subsets. Frequency, GFP^+^, dtTomato^+^ and total cell counts were calculated, exported and visualized using Prism (v9.0) (GraphPad).

### RNA Isolation and RT-qPCR

RNA was isolated using RNAsolv (Omega Bio-tek) or NucleoSpin RNA XS Kit (Macherey-Nagel) according to the manufacturer’s protocol. For SYBRGreen-based RT-qPCR, cDNA was generated from RNA using a high-capacity cDNA Reverse Transcription Kit (Applied Biosystems). Volcano3G® RT-PCR Probe 2x Master Mix (myPOLS)-based RT-qPCR and SYBRGreen-based qPCR was performed using a StepOnePlus Real-Time PCR System/QuantStudio™ 3 System (Applied Biosystems). All genes were normalized to *Actb* (β-actin). Primers are listed in table EV1.

### Genomic DNA Isolation and PCR

Genomic DNA from isolated thymocytes was purified using the SV Total Genomic DNA Isolation Kit (Promega) according to the manufacturer’s protocol. Amplification of the floxed *Nr5a2* gene, the Cre-deleted *Nr5a2* gene and myogenin as a loading control was performed by PCR (95°C, 2 min; 35 cycles: 95°C, 57°C 30 sec each step, 68°C 40 sec; 68°C 5 min). Primers for *loxP* site, the deleted *Nr5a2* gene and myogenin are listed in table EV2. PCR products were then separated by 2% agarose gel electrophoresis and visualized using ImageQuantLAS400 (GE Healthcare Life Sciences).

### Hematoxylin and Eosin Staining

Formalin-fixed and paraffin-embedded murine thymus and spleen were deparaffinized and dehydrated prior to hematoxylin and eosin (H&E) staining.

### Enzyme-linked Immunosorbent Assay (ELISA)

Secretion of IL–2, IL-4, IL-6, Tumor Necrosis Factor (TNF) and Interferon (IFN)-γ from splenocytes after T cell receptor activation with 3 µg/ml plate bound anti-CD3ε antibody and 1 µg/ml soluble anti-CD28 antibody for 48 h, as well as restimulation with 1 ng/ml PMA and 200 ng/ml ionomycin at 42 h post-stimulation for additional 6 h, was quantified by ELISA according to the manufacturer’s protocol (BioLegend). Additionally, serum TNF and IFN-γ of Rag2^-/-^ mice from colitis experiments were measured by ELISA following the same protocol. Antibodies used are listed in RTT.

### Seahorse Basal ATP Rate

Glycolytic and mitochondrial ATP production rates of thymocytes or splenic CD4^+^ and CD8^+^ T cells were determined according to the manufacturer’s protocol (Agilent). Isolated cells were resuspended in Seahorse XF Dulbecco’s Modified Eagle Medium (DMEM) medium pH 7.4 supplemented with 18 mM glucose, 1 mM pyruvate, and 2 mM L-glutamine. Before calibration of the pre-hydrated XFe24 sensor cartridge in the Flux Analyzer XFe24, port A was filled with 1.5 µM oligomycin, port B with 1 µM carbonyl cyanide-p-trifluoromethoxyphenylhydrazone and port C with a mixture of 0.5 µM rotenone, 0.5 µM antimycin A and 1 µg/ml Hoechst. For normalization, cell numbers were determined based on an automated algorithm in Cellomics ArrayScanTM VTI High-Content Screening (CellomicsTM) by counting Hoechst^+^ nuclei. ATP production rate was analyzed using Agilent Seahorse Analytics Online software and normalized to control for thymocyte measurement and unstimulated control for splenic T cell measurement.

### Statistical Analyses

Statistical analyses were performed with GraphPad Prism (v9.0). Data were tested for normal distribution using the D’Agostino-Pearson omnibus, Shapiro-Wilk, or Kolmogorov-Smirnov test. The alpha level of significance was set at 0.05, with *p < 0.05, **p < 0.01, ***p < 0.001, and ****p < 0.001 for all tests. One-way ordinary ANOVA with Dunnett’s multiple comparison test, ordinary two-way ANOVA with Turkey multiple comparison test, unpaired Student’s T-test and multiple unpaired Student’s t-test with Holm-Šídák multiple comparison test were performed. Details of each statistical test performed on the data are provided in the corresponding figure legend. Unless otherwise noted, data are presented as mean ± SEM.

## Authors’ contributions

AW and TB conceived the study. AW performed the experiments, designed and analyzed the data. NK contributed to the establishment of the *ex vivo* LRH-1 inducible knockout system in thymocytes. RHME, LD and FR helped with the colitis experiments. AW, VMM and LD performed histological analyses. All authors have discussed the results. AW wrote the manuscript, and VMM and DFL helped with the manuscript design. TB edited the manuscript and acquired funding.

## Acknowledgements

The authors thank the Animal Facility (TFA) and Flow Cytometry Core Facility (FlowKon) of the University of Konstanz for technical support, Daniela Finke (University of Basel) for providing the OP9-DL1 cells, Kristina Schoonjans (EPF Lausanne) for the LRH-1^L2/L2^ mice, Freddy Radke (EPFL Lausanne) for the Lck-Cre mTmG mice, and Michael Scharl and Marianne Spalinger (University Hospital Zurich) for the Rag2^-/-^ mice. This study was supported by a grant from the German Research Foundation (DFG) to TB (BR 3349/4-2). RHME is supported by the German Academic Exchange Service (DAAD) under the Research Grants - Doctoral Programmes in Germany, 2023/24 (57645448). DFL is supported by the Irène and Max Gsell Foundation.

## Disclosure and competing interest statement

All authors declare no competing interests.

## Abbreviations

4-HT: 4-hydroxytamoxifen
BSA: bovine serum albumin
BrdU: bromodeoxyuridine
CD: cluster of differentiation
cKO: conditional knockout
DAPI: 4′,6-diamidino-2-phenylindole
DN: double negative
DP: double positive
DTT: dithiothreitol
EDTA: ethylenediaminetetraacetic acid
FBS: fetal bovine serum
HBSS: hanks balanced salt solution
H&E: hematoxylin and eosin
HEPES: 4-(2-hydroxyethyl)-1-piperazineethanesulfonic acid
HS: horse serum
IEL: intraepithelial lymphocytes
IFN: interferon
i.p.: intraperitoneal
ILC: innate lymphoid cells
iKO: inducible knockout
iNK: immature natural killer cells
LP: lamina propria
LPL: lamina propria lymphocytes
LRH-1: Liver Receptor Homolog 1 (Nr5a2)
mNK: mature natural killer cells
mLN: mesenteric lymph nodes
mIL: murine interleukin
mTmG: membrane-targeted tandem dimer Tomato membrane-targeted GFP
NK: natural killer cells
NKT: natural killer T cell-like cells
PMA: phorbol-12-myristat-13-acetate
SP: single positive
TCR: T cell receptor
T_Eff_: effector T cells
T_reg_: regulatory T cells
TNF: tumor necrosis factor
UMAP: Uniform Manifold Approximation and Projection

## Supplementary Figure Legends

**Supplementary Figure 1:**
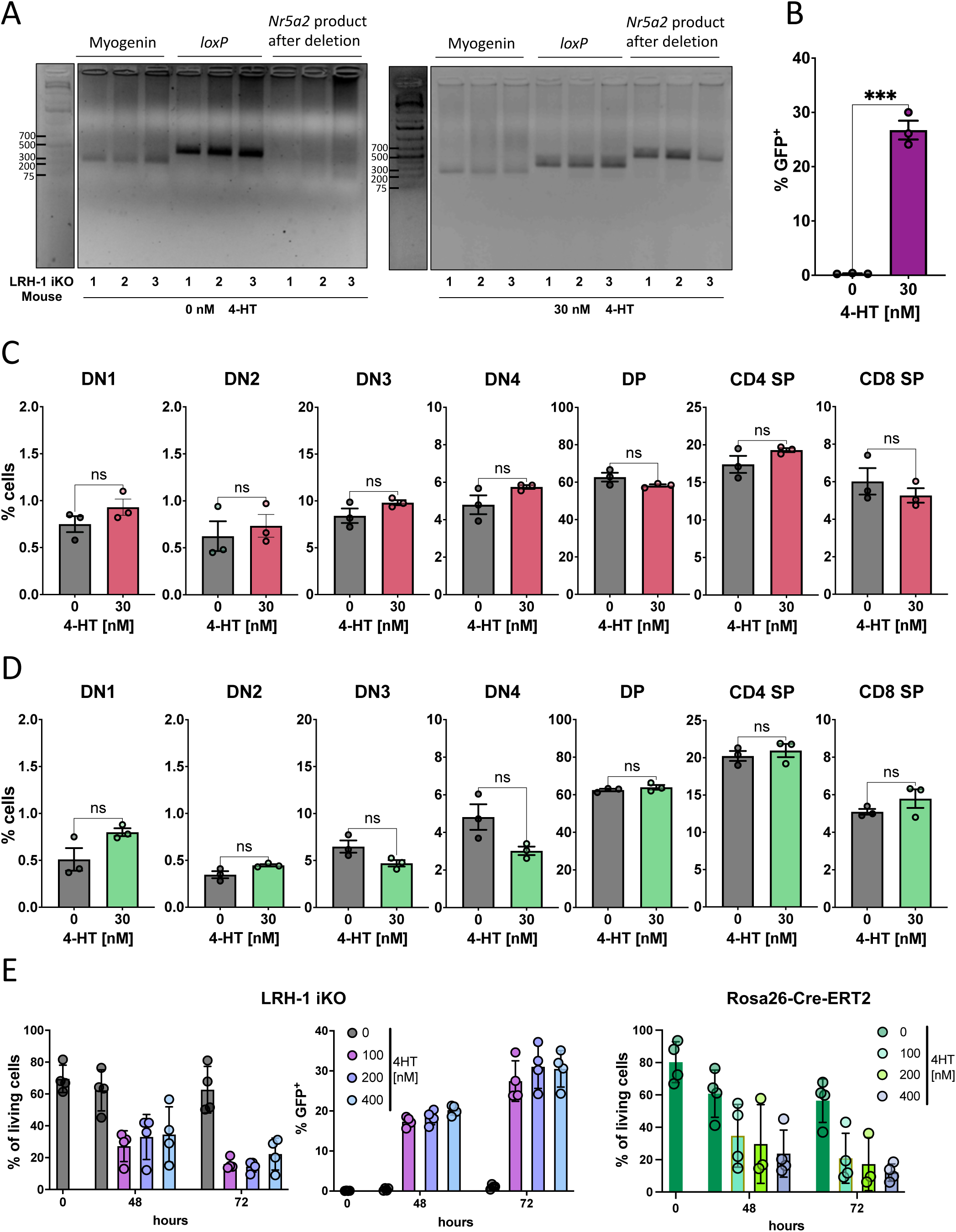
Validation of an *ex vivo* 4-hydroxytamoxifen-inducible LRH-1 knockout system in thymocytes. (A) Agarose gel electrophoresis of LRH-1 iKO thymocytes, untreated (0 nM 4-HT) or treated with 30 nM 4-HT for 20 h to confirm LRH-1 deletion after 4-HT addition by PCR. *Nr5a2* loxP site (380 bp) and *Nr5a2* gene product after deletion (525 bp) were detected. *Myog* (myogenin) (250 bp) was used as a loading control (n = 3 mice). Flow cytometry analysis of GFP-positive live LRH-1 iKO thymocytes (B) and frequency of live thymocyte stages of control LRH-1^L2/L2^ (C) and control Rosa26-Cre-ERT2 mice (D), untreated or treated twice with 30 nM 4-HT (0 h, 24 h) and analyzed after 72 h. Bars show mean ± SEM, dots represent individual mice (n = 3 per group). (E) Flow cytometry analysis of frequency of live thymocyte stages in LRH-1 iKO and Rosa26-Cre-ERT2 control thymocytes, and GFP-positive live LRH-1 iKO thymocytes. Thymocytes from LRH-1 iKO or Rosa26-Cre-ERT2 control mice were left untreated or treated with the indicated concentration of 4-HT, and measured after 0 h, 48 h and 72 h. Bars show mean ± SEM, dots represent individual mice (n = 4 per group). Thymocytes were co-cultured with OP9-DL1 cells and treated with IL-7 (1 ng/ml). Statistical significance was determined by unpaired Student’s t-test, ns: not significant, *** p<0.001. DN = double negative, DP = double positive, SP = single positive.

**Supplementary Figure 2:**
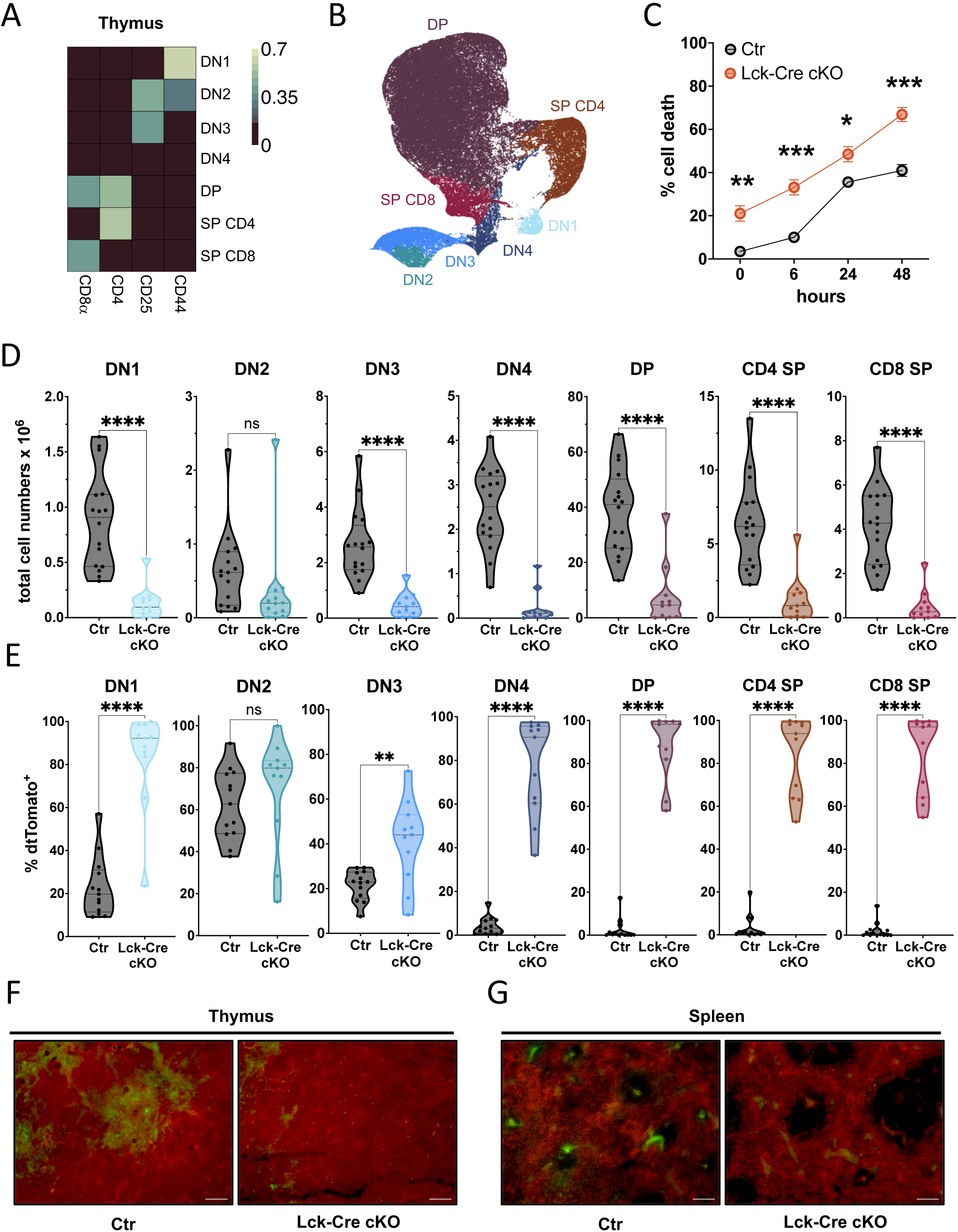
Characterization of thymocytes from Lck-Cre LRH-1 cKO mice. (A) Thymocyte surface marker expression heatmap of CD8α, CD4, CD25 and CD44 expression according to FlowSOM clustering of double negative (DN)1 (CD44^+^), DN2 (CD44^+^CD24^+^), DN3 (CD25^+^), double positive (DP; CD4^+^CD8^+^), single positive (SP) CD4 (CD4^+^), SP CD8 (CD8^+^) subsets, and (B) UMAP clusters from high dimensional flow cytometry analysis of Lck-Cre cKO and Ctr thymocytes. Data were pre-gated for live cells. (C) Flow cytometry analysis of DAPI-positive (dead) thymocytes from Ctr and Lck-Cre cKO mice co-cultured with OP9-DL-1 cells and treated with IL-7 (1 ng/ml) for the indicated time points. Points represent mean ± SEM (n = 5 mice per group). High-dimensional flow cytometry analysis of thymocyte numbers (D) and dtTomato-positive (E) live thymocyte stages from Ctr and Lck-Cre cKO mice. Each point in the violin plots represents an individual mouse (n = 16 Ctr, n = 11 Lck-Cre cKO). (F) Representative fluorescence images of thymus and spleen sections, showing dtTomato (red) and GFP (green) fluorescence, from 13-week-old Lck-Cre cKO and Ctr mice. Scale bar = 100 µm. Statistical significance was performed using multiple unpaired Student’s t-test with Holm-Šídák multiple comparison test (C) and unpaired Student’s t-test (D, E), ns: not significant, * p<0.05, ** p<0.005, **** p<0.0001. DN = double negative, DP = double positive, SP = single positive.

**Supplementary Figure 3:**
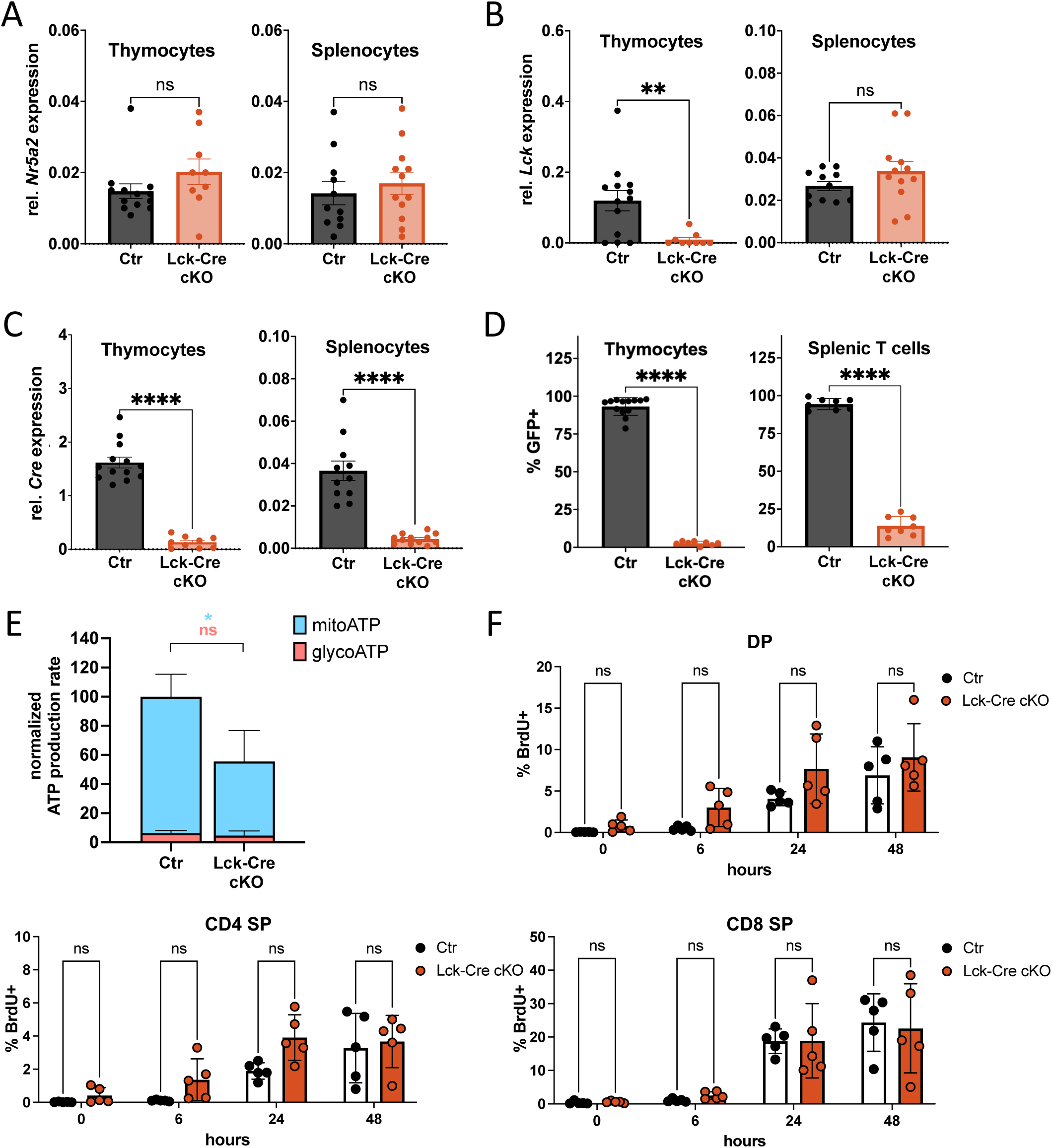
Characterization of escaping LRH-1 cKO thymocytes and splenocytes. Relative RT-qPCR of *Nr5a2* (A), *Lck* (B) and *Cre* (C) from isolated Lck-Cre cKO thymocytes and splenocytes. Gene expression was normalized to *Actb*. Bars show mean ± SEM, dots represent individual mice (n= 12 Ctr, n= 9-11 Lck-Cre cKO). (D) Flow cytometry analysis of GFP-positive live thymocytes and splenic T cells isolated from Ctr and Lck-Cre cKO mice. Bars show mean ± SEM, dots represent individual mice (n= 8-13 Ctr, n= 8 Lck-Cre cKO). (E) Seahorse basal ATP rate assay of isolated thymocytes. Bar chart represents mean ± SEM, ATP production rate normalized to CTR (n = 3-4 mice per group). (F) Flow cytometry analysis of BrdU-positive live DP, CD4 SP and CD8 SP thymocytes from Ctr and Lck-Cre cKO mice at indicated time points. Bars show mean ± SEM, dots represent individual mice (n = 5 per group). Thymocytes were co-cultured with OP-DL1 cells and treated with IL-7 (1 ng/ml). Statistical significance was performed using unpaired Student’s t-test (A-E) with comparison of mitoATP and glycoATP independently (E), and multiple unpaired Student’s t-test with Holm-Šídák multiple comparison test (F), ns: not significant, * p<0.05, ** p<0.005, **** p<0.0001. DP = double positive, SP = single positive.

**Supplementary Figure 4:**
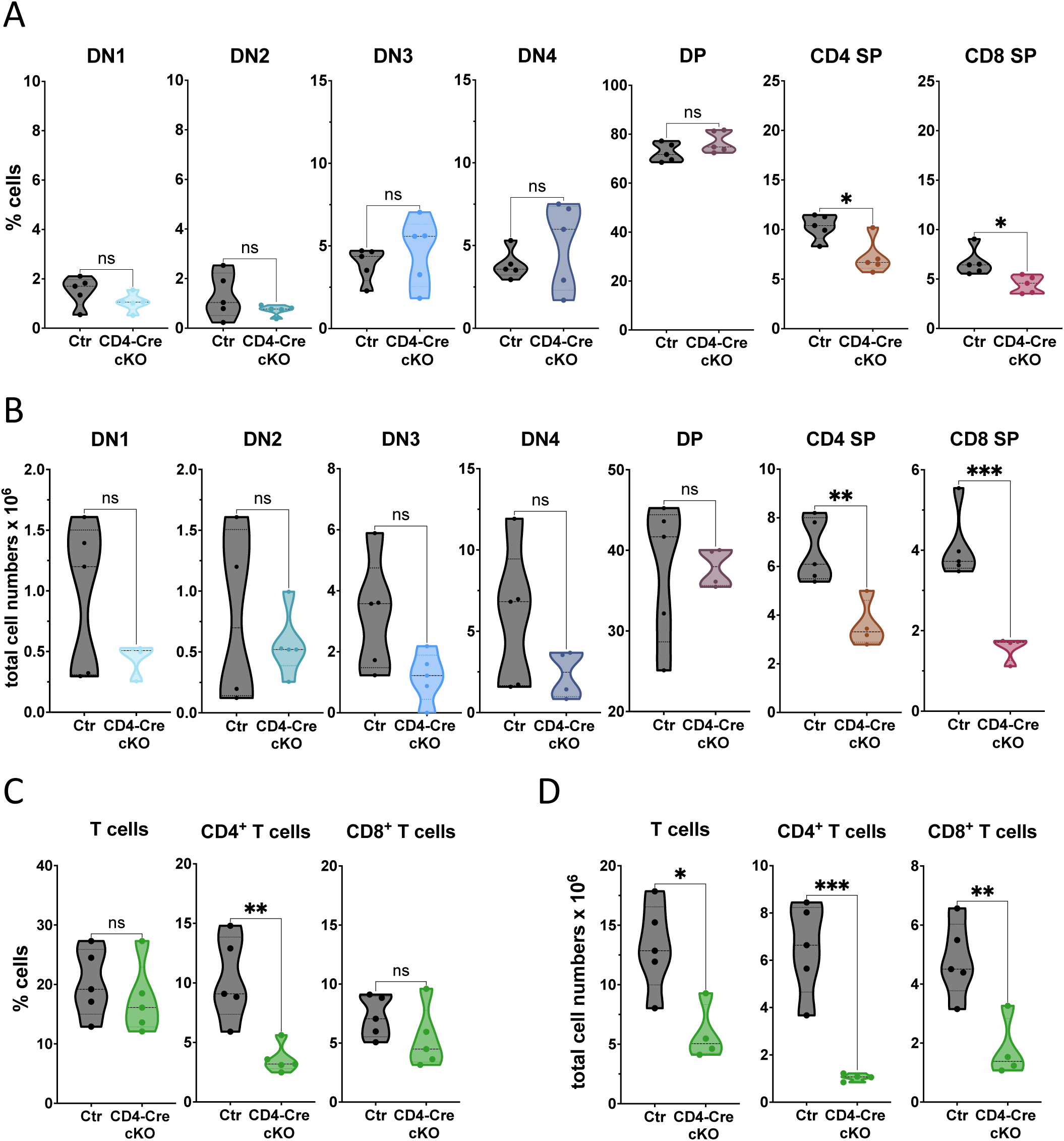
T cell distribution in CD4-Cre LRH-1 cKO mice is altered only in single positive thymocytes and mature CD4^+^ T cells. Flow cytometry analysis of thymocyte frequency (A) and total cell count (B) of different thymocyte stages in CD4-Cre (Ctr) and CD4-Cre x LRH-1 (CD4-Cre cKO) mice. Flow cytometry analysis of isolated splenic T cell frequency (C) and total cell count (D) in Ctr and CD4-Cre cKO mice. Dots represent individual mice (n = 4 per group). Statistical analysis was performed using unpaired Student’s t-test, ns: not significant, * p<0.05, ** p<0.005, *** p<0.001. DP = double positive, SP = single positive.

**Supplementary Figure 5:**
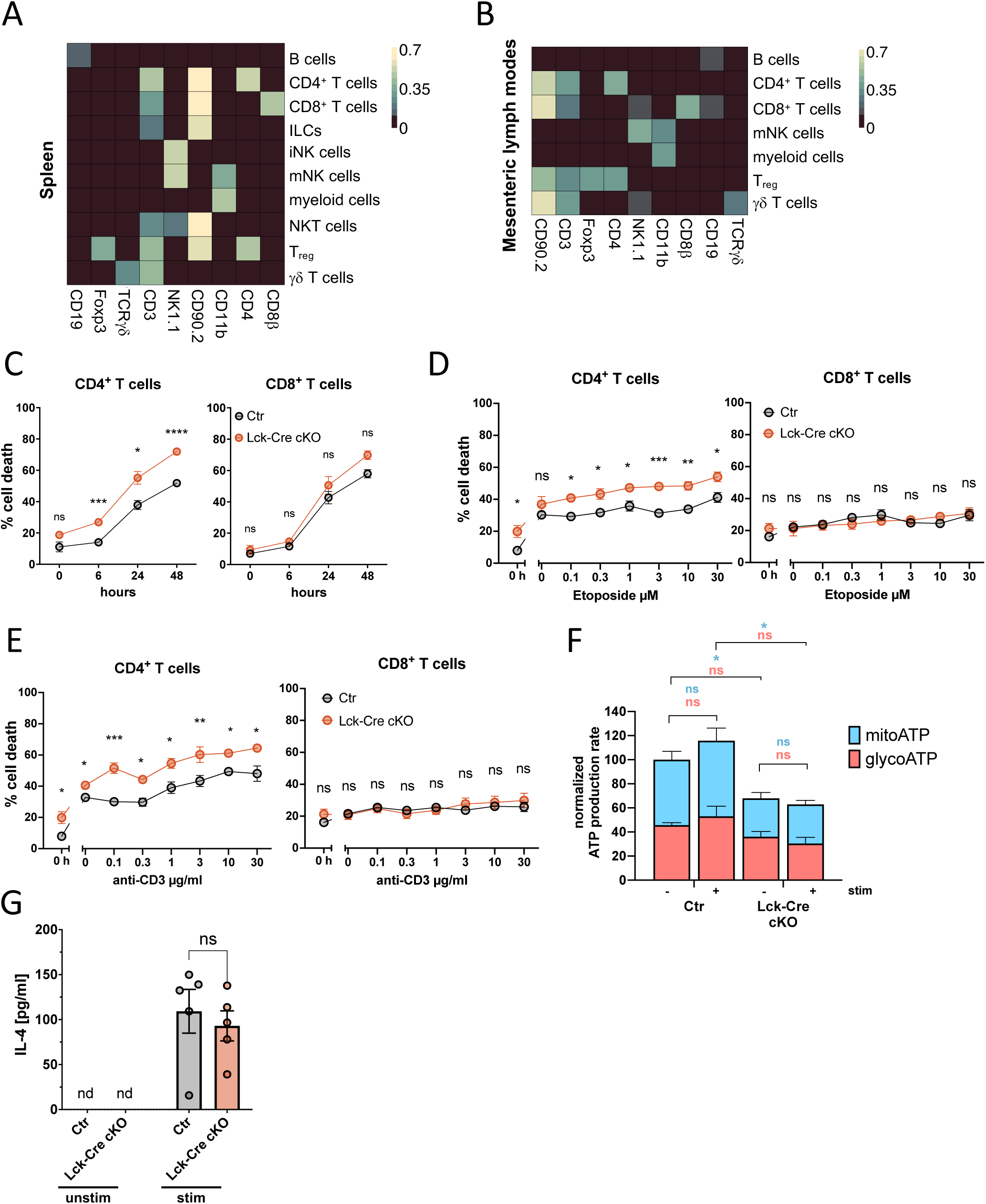
Characterization of T cells from spleen and mesenteric lymph nodes of Lck-Cre control and Lck-Cre LRH-1 cKO mice. Surface marker expression heatmap of cells isolated from spleen (A) and mesenteric lymph nodes (B). CD19, Foxp3, TCRγδ, CD3, NK1.1, CD90.2, CD11b, CD4, CD8β expression according to FlowSOM clustering. Data were pre-gated to live CD45^+^ cells. B cells (CD19^+^), CD4^+^ T cells (CD4^+^ CD90.2^+^ CD3^+^), CD8^+^ T cells (CD8β^+^ CD90.2^+^ CD3^+^), innate lymphoid cells (ILCs; CD9.2^+^ CD3^+^), myeloid cells (CD11b^+^), immature natural killer cells (iNK; NK1.1^+^), mature NK cells (mNK; NK1.1^+^ CD11b^+^), NKT cells (NK T cell-like cells; NK1.1^+^ CD90.2^+^), regulatory T cells (T_reg_; CD4^+^ CD90.2^+^ CD3^+^ Foxp3^+^), gamma-delta T cells (γδ T cells; CD3^+^ CD90.2^+/-^ TCRγδ^+^). (C) Flow cytometry analysis of cell death (DAPI^+^) in CD4^+^ and CD8^+^ T cells from Ctr and Lck-Cre cKO mice measured at the indicated time points. Points represent mean ± SEM (n = 5 mice per group). Flow cytometry analysis of cell death (DAPI^+^) in CD4^+^ and CD8^+^ T cells after etoposide treatment (D) (6 h) and plate-bound anti-CD3ε stimulation (E) (24 h). Dots represent mean ± SEM (n = 3-4 mice). (F) Seahorse basal ATP rate assay of splenic CD4^+^ and CD8^+^ T cells stimulated with plate-bound anti-CD3ε (3 µg/ml) and anti-CD28 (1 µg/ml), or left unstimulated for 24 h. Bar blot represents mean ± SEM. ATP production rate was normalized to unstimulated controls (n = 3 mice per group). (G) IL-4 secretion in splenocytes. T cells were primed with plate bound anti-CD3ε (3 µg/ml) and anti-CD28 (1 µg/ml), or left unstimulated for 42 h. Cells were then restimulated with PMA (1 ng/ml) and ionomycin (200 ng/ml) for an additional 6 h. Points represent mean ± SEM (n = 5 mice). Statistical significance was analyzed using multiple unpaired Student’s t-test with Holm-Šídák multiple comparison test (C-E) and ordinary two-way ANOVA with Turkey multiple comparison test (F, G) comparing mitoATP and glycoATP independently (F). nd = not detected, ns: not significant, * p<0.05, ** p<0.005, *** p<0.001, **** p<0.0001. Unstim = unstimulated, stim = stimulated.

**Supplementary Figure 6:**
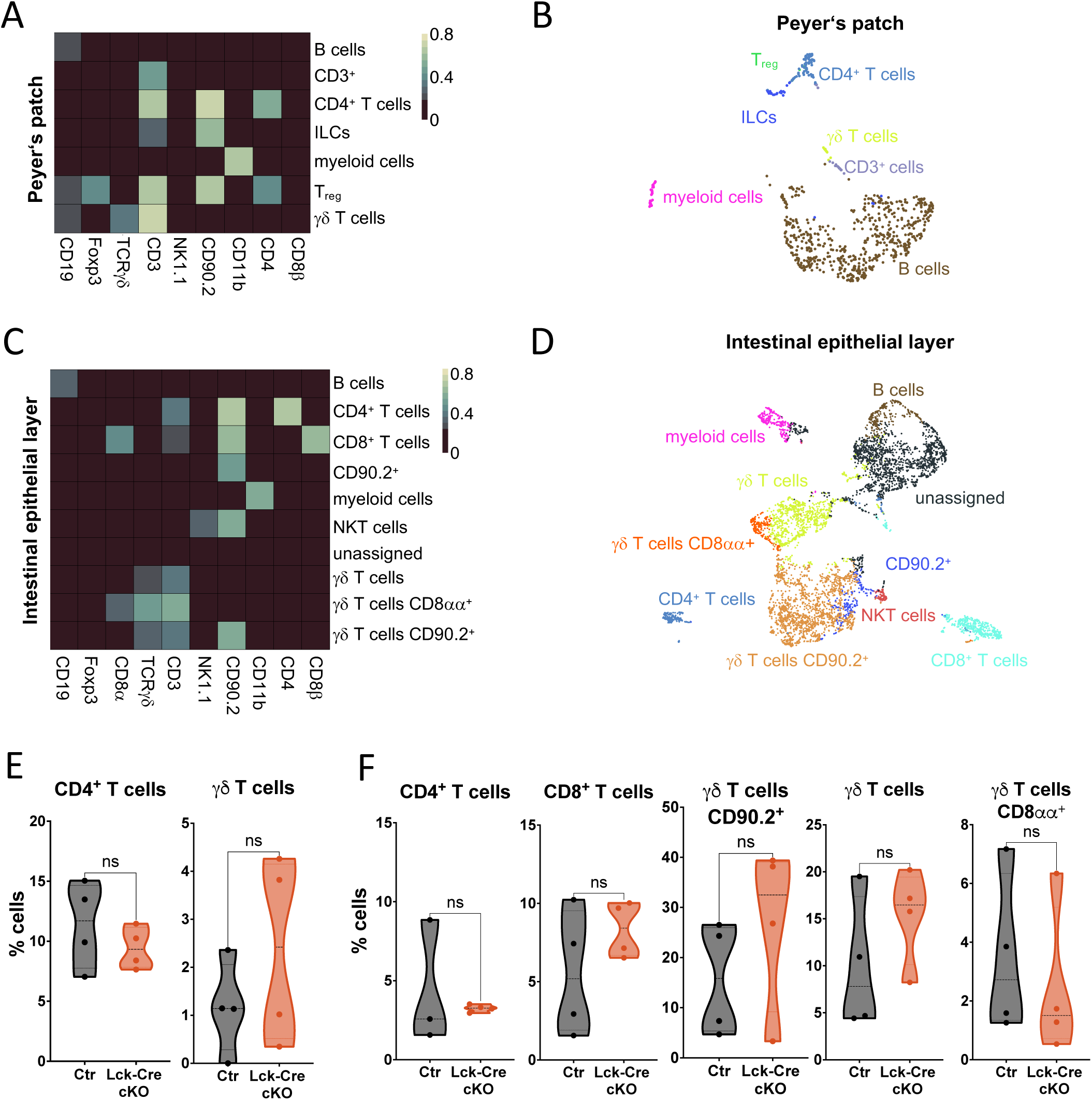
Characterization of T cells from Peyer’s patches and intestinal epithelial layer in Lck-Cre control and Lck-Cre LRH-1 cKO mice. Surface marker expression heatmap of cells isolated from Peyer’s patches (A) and intestinal epithelial layer (C). Expression of CD19, Foxp3, TCRγδ, CD3, NK1.1, CD90.2, CD11b, CD4, CD8α and CD8β in intestinal immune cells according to FlowSOM clustering. UMAP clusters of cells from Peyer’s patches (B) and intestinal epithelial layer (D). Data were pre-gated to live CD45^+^ cells. B cells (CD19^+^), CD4^+^ T cells (CD4^+^ CD90.2^+^ CD3^+^), CD8^+^ T cells (CD8β^+^ CD8α^+^ CD90.2^+^ CD3^+^), CD3^+^, CD90.2^+^, myeloid cells (CD11b^+^), innate lymphoid cells (ILC; CD9.2^+^ CD3^+^), natural killer T cell-like cells (NKT cells; NK1.1^+^ CD90.2^+^), regulatory T cells (T_reg_; CD4^+^ CD90.2^+^ CD3^+^ Foxp3^+^), gamma-delta T cells (γδ T cells; CD3^+^ CD90.2^+/-^TCRγδ^+^ CD8αα^+/-^) are shown. High-dimensional flow cytometry analysis of the frequency of T cells from Peyer’s patches (E) and intestinal epithelial layer (F) from Ctr and Lck-Cre cKO mice. Each point in the violin plot represents individual mice (n = 4 per group). Statistical significance was analyzed by unpaired Student’s T-test. ns: not significant.

**Supplementary Figure 7:**
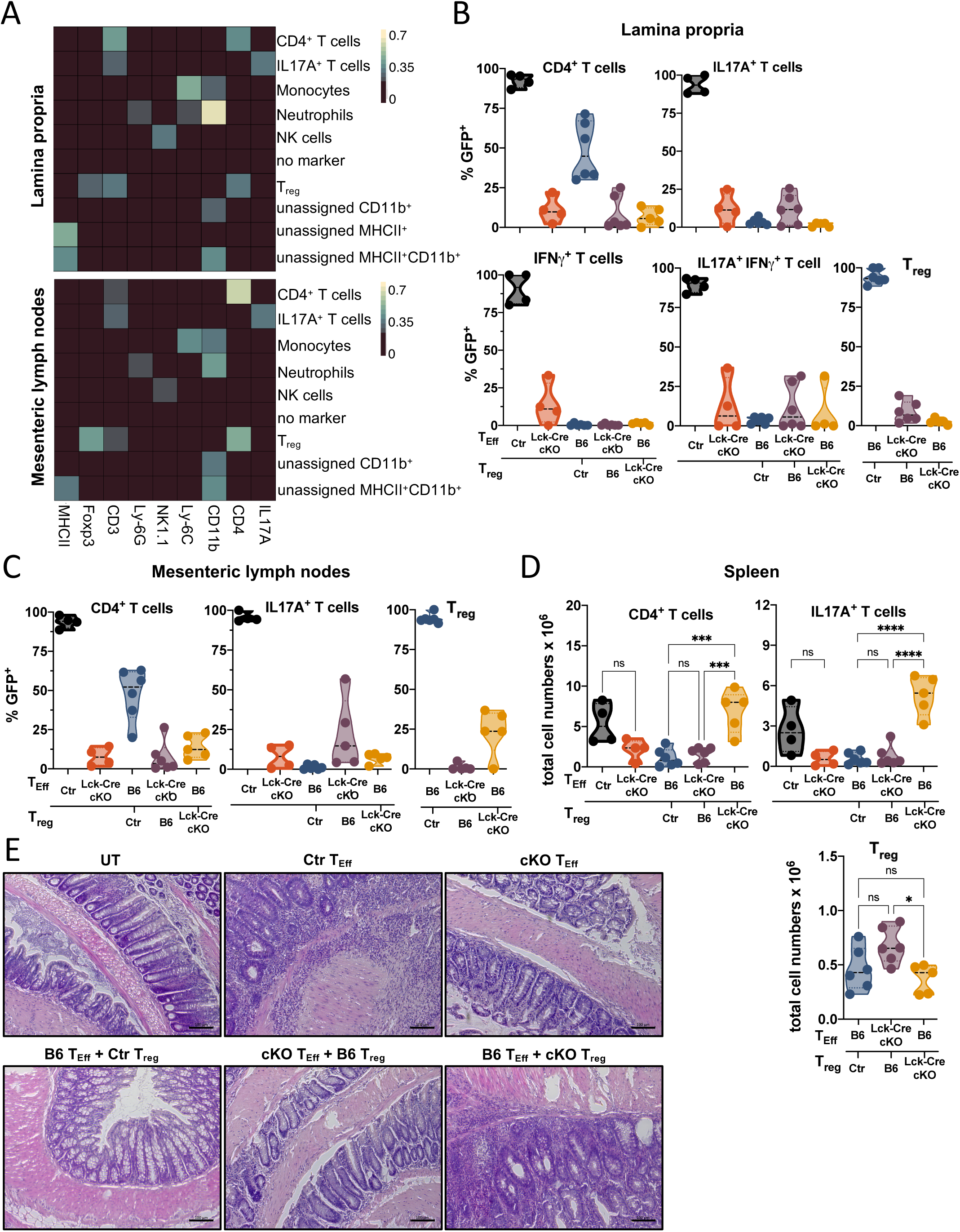
Immunophenotyping of immune cells from the lamina propria, mesenteric lymph nodes and spleen during effector T cell-mediated colitis and regulatory T cell-mediated protection. (A) Surface marker expression heatmap of immune cells isolated from colonic lamina propria and mesenteric lymph nodes. Expression of MHCII, CD3, Ly-6G, NK1.1, Ly-6C, CD11b, CD4, IL17A and Foxp3 according to FlowSOM clustering were used to define the immune cell subsets. High-dimensional flow cytometry analysis of GFP-positive T cells from the colonic lamina propria (B), the mesenteric lymph nodes (C) and total T cell numbers from the spleen (D) of Rag2^-/-^ mice. Data were pre-gated to live CD45^+^ cells. Each point in the violin plots represents an individual mouse (n = 4 T_Eff_ transfer of Ctr and Lck-Cre cKO, n = 5-6 T_Eff_ and T_reg_ co-transfer of Ctr, B6 and Lck-Cre cKO). (E) Representative H&E-stained colon sections from untreated (UT) Rag2^-/-^ and Rag2^-/-^ mice receiving T_Eff_ from the Ctr and Lck-Cre cKO (top panel) or co-transfers of T_Eff_ from B6 with T_reg_ from Ctr or Lck-Cre cKO or T_Eff_ from Lck-Cre cKO with T_reg_ from B6 (bottom panel). Scale bar = 100 µm. Statistical significance was determined by one-way ordinary ANOVA with Dunnett’s multiple comparison test. ns: not significant, * p<0.05, *** p<0.001, **** p<0.0001.

**Expanded View Table 1.**
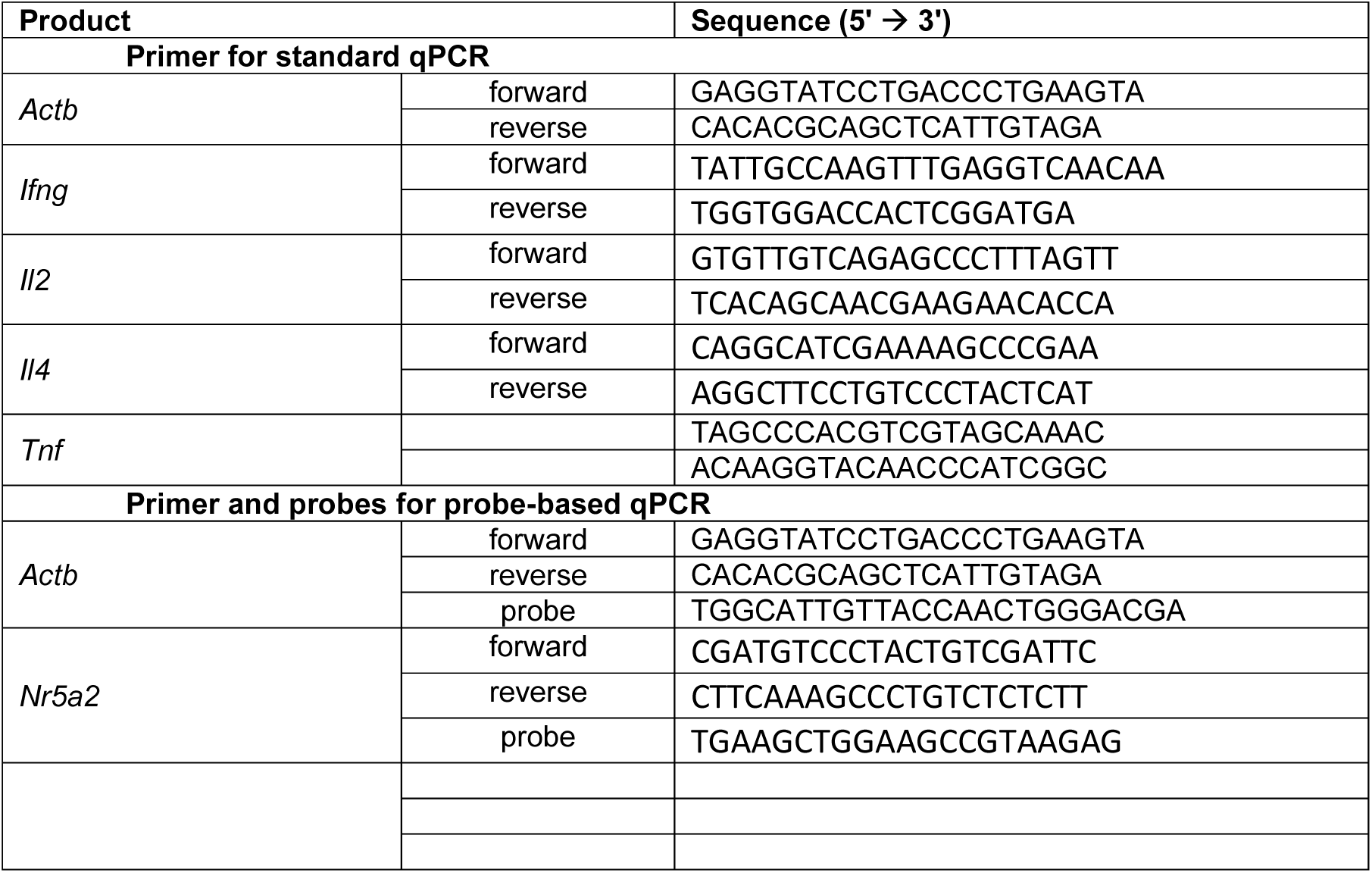
(related to RTT) Primers sequences for quantitative real-time PCR (qPCR)

**Expanded View Table 2.**
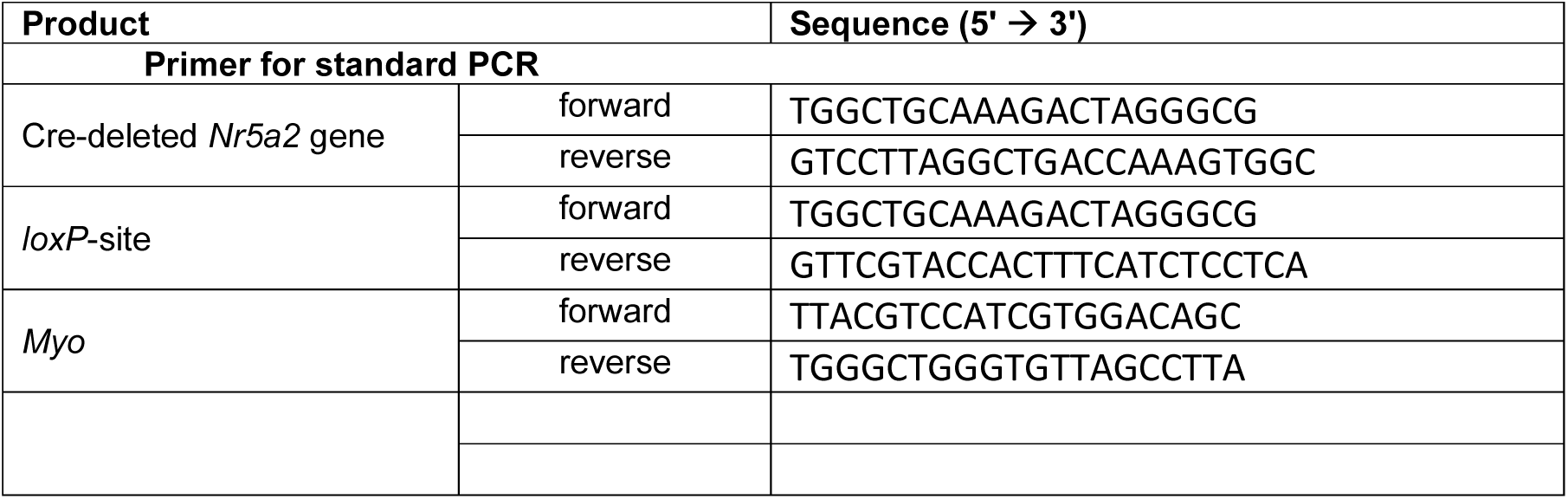
(related to RTT) Primers sequences for real-time PCR (PCR)

## Notes

### Competing Interest Statement

The authors have declared no competing interest.

